# Macrophage efferocytosis is controlled by epigenetic modifications mediated by RBPJ

**DOI:** 10.1101/2025.10.10.681723

**Authors:** Samreen Sadaf, Xinyi Zhang, Liqun Lei, Jixing Shen, Imran Jamal, Ganesh Modugu, Partha Dutta

**Author notes:** Corresponding author: **Partha Dutta, DVM, PhD**, Associate Professor of Medicine and Immunology, Center for Pulmonary Vascular Biology and Medicine, Pittsburgh Heart, Lung, Blood, and Vascular Medicine Institute, University of Pittsburgh School of Medicine, 200 Lothrop Street BST1720.1, Pittsburgh, PA USA 15213. These authors equally contributed to this work (Samreen Sadaf, Xinyi Zhang). **Declaration of interest:** The authors have declared that no conflict of interest exists.

## Abstract

Efferocytosis, phagocytic clearance of apoptotic cells, is crucial for inflammation resolution and maintenance of tissue homeostasis. However, it is not known how epigenetic alterations govern macrophage-mediated efferocytosis. A Cleavage Under Targets and Release Using Nuclease (CUT&RUN) sequencing revealed a genome-wide selective suppression of H3K9me3, a heterochromatin mark that represses gene activity, in macrophages undergoing efferocytosis. Moreover, Recombination Signal Binding Protein for Immunoglobulin Kappa J region (RBPJ), which is a transcription factor typically involved in the canonical Notch signaling process, dampened this epigenetic modification, enhanced apoptotic cell clearance, and suppressed inflammation by mouse atherosclerotic plaque, alveolar, peritoneal, and bone marrow-derived macrophages and human primary macrophages. Inhibition of the Notch signaling in macrophages significantly reduced efferocytosis whereas activation of this signaling augmented apoptotic debris clearance. Mechanistically, RBPJ upregulated *Stard13* and *Arsg* by diminishing H3K9me3 on their promoters. *Stard13* promoted efferocytosis by magnifying actin polymerization via inhibition and activation of Rho and RAC GTPases, respectively. Genetic and pharmacological inhibitions of SUV39H1/H2, the methyltransferases that are responsible for H3K9 trimethylation, amplified the expression of *Stard13* and *Arsg*, and augmented efferocytosis by *RBPJ*^-/-^ macrophages. In sum, this study shows epigenetic regulation of efferocytosis in tissue macrophages.

**Graphical Abstract:** 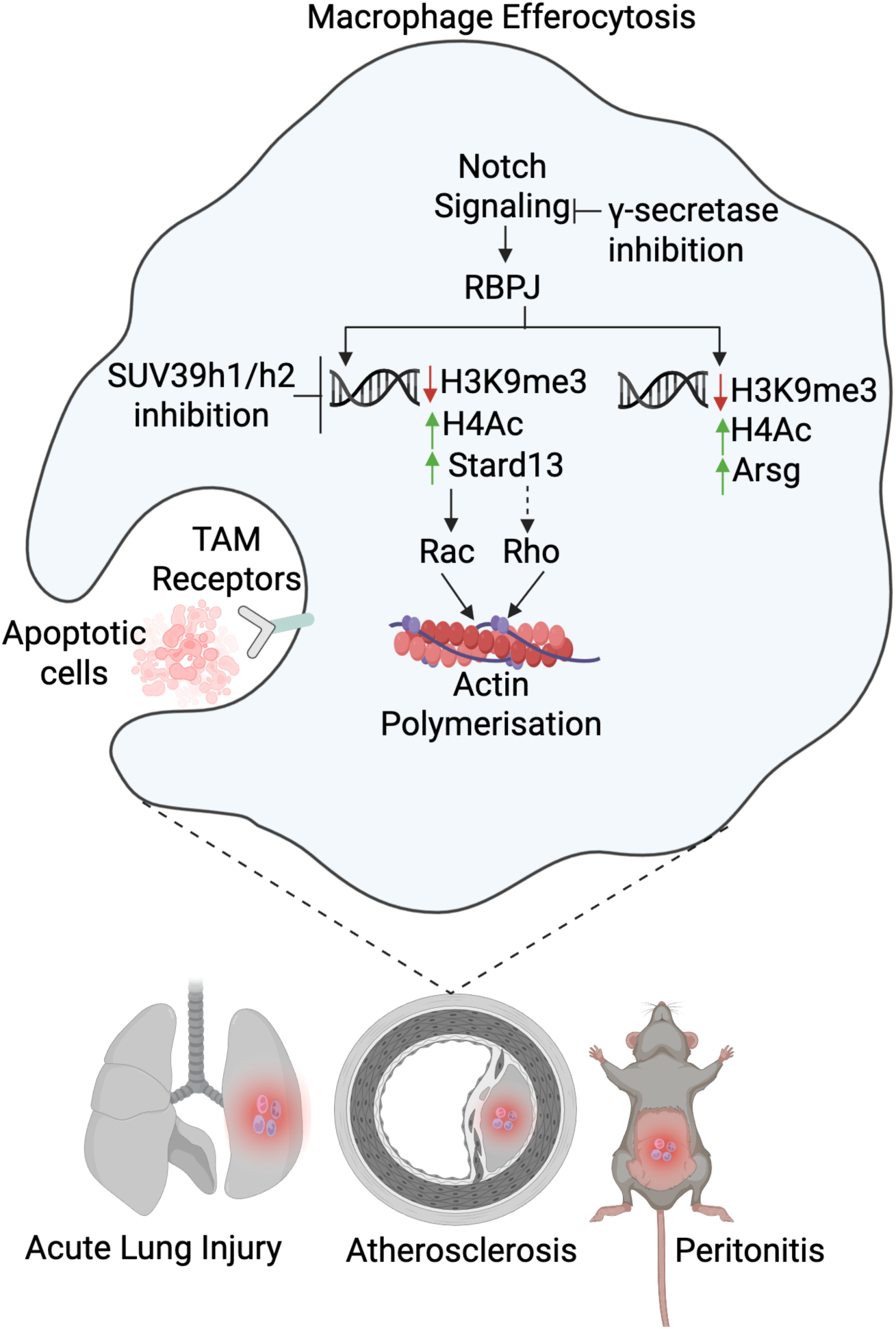

## Introduction

Efferocytosis, the process by which phagocytes eliminate apoptotic cells, is essential in preventing the necrosis of dead cells and inducing anti-inflammatory and pro-resolving responses in phagocytes like macrophages(1). There are various stages of efferocytosis, such as the (i) sensing phase, during which phagocytes and apoptotic cells converse via mediators; (ii) contact phase, during which apoptotic cells and phagocytes establish ligand-receptor interactions; and (iii) digestion phase, during which apoptotic bodies are internalized and processed (2). Studies have revealed that certain diseases are characterized by impaired macrophage-mediated efferocytosis, such as acute peritonitis, lung injury, and atherosclerosis, promoting disease progression(2, 3). However, the mechanisms of defective efferocytosis in disease conditions are not clear.

Epigenetic modifications of genes by histone-modifying enzymes, such as methyltransferases and acetyltransferases, in immune cells alter their functions in different diseases including cancer(4), pneumonia(5), diastolic dysfunction(6), and non-alcoholic steatohepatitis(7). For example, histone methyltransferases decreased anti-tumor ability (8) and increased the expression of skin-homing receptors by dampening Stat5 phosphorylation(9) of immune cells. Additionally, Dot1l, an H3K79me2/3 methyltransferase, heightened IL-6 and IFN-β production(10), and prevented antigen-independent differentiation of CD8 T cells(11). Methyltransferases also promoted transition of pro-inflammatory to reparative macrophages in diabetic wounds(12). Similarly, acetyl transferases can modulate inflammation. Histone acetyl transferase p300 has been shown to catalyze H3K27 acetylation to increase IL-9 production(13). Histone demethylases, which removes methyl groups from histones, can affect macrophage differentiation and function(14). For example, Kdm4a encouraged M1 macrophage polarization(15), Kdm6a, an H3K27me2/3 demethylase, promoted IL-6 and IFN-β production in macrophages(16), and Kdm5b induced immune evasion by recruiting H3K9 methyltransferases (8). Epigenetic readers, such as BRD9, can modulate macrophage inflammatory response by augmenting glucocorticoid receptor activity(17). Furthermore, recent research has revealed that epigenetic changes like DNA methylation, histone modifications, and RNA regulation play a major role in macrophage polarization and differential activation, making them viable therapeutic targets for the management of cardiovascular disease (18). As such, there is a consensus that epigenetic modifications can alter macrophage-mediated inflammation. However, if apoptotic cell clearance by macrophages is regulated by epigenetic alterations is not known.

A highly conserved DNA-binding protein known as the recombination signal binding protein for immunoglobulin kappa J region (RBPJ) is essential for determining the fate of cells(19–21). It is ubiquitously expressed and can mediate transcriptional activation and repression of Notch target genes by interacting with coactivators or corepressors, respectively(19, 22, 23). Studies have reported that RBPJ involves in the regulation of both lymphoid and myeloid cells(24–29). For example, endothelial RBPJ signaling can promote inflammatory leucocyte recruitment in atherosclerosis(30). Besides, the transcription factor also mediates macrophage polarization via stimulating several downstream genes(25–28). The connection of RBPJ with macrophage raised the possibility that RBPJ does not simply modulate the polarization of macrophage but might also influence its function, such as efferocytosis, a hypothesis that has not been addressed thus far. Moreover, our RNA sequencing analysis comparing monocyte-derived and tissue resident macrophages revealed that the expression of the genes regulated by RBPJ was significantly altered in monocyte-derived macrophages highlighting the importance of this transcription factor in inflammation. To this end, we induced atherosclerosis in myeloid-specific *Rbpj*-deficient mice. To our surprise, we observed larger necrotic core area in atherosclerotic plaques in these mice compared to age-matched control mice. Since impaired efferocytosis in atherosclerosis is proposed to mediate necrotic core enlargement, we sought to delineate the merit of macrophage RBPJ in efferocytosis.

In this study, we had several goals. First, we aimed to determine how epigenetic changes affected macrophage-mediated efferocytosis. Second, we wanted to understand the contribution of RBPJ in this process. To understand the function of macrophage RBPJ in inflammation resolution, we used three mouse models. 1. lipopolysaccharide (LPS)-induced lung damage is a model of acute lung injury (ALI). It has been reported that defects in efferocytosis by alveolar macrophages can cause persistent pneumonia while efficient efferocytosis can prevent lethal lung damage from lung infection (31, 32). 2. Zymosan-induced peritonitis is widely used as an acute inflammation followed by its resolution. In this inflammatory setting, macrophages are known to proliferate and clear apoptotic neutrophils to promote inflammation resolution (33–35). 3. Atherosclerosis is a disease of blood vessels wherein macrophage-mediated efferocytosis is compromised, resulting in the accumulation of necrotic debris in atherosclerotic plaques and their rupture ensuing blockade of blood flow to vital organs (36). We showed that RBPJ enhanced macrophage efferocytosis. This transcription factor in macrophages promoted efferocytosis by inhibiting H3K9me3, an epigenetic mark that suppresses gene expression, on the promoters of the RBPJ-dependent genes, such as *Stard13* and *Arsg*, resulting in their heightened expression. Stard13 inhibited and activated Rho and RAC GTPases, respectively, promoting actin polymerization and engulfment of apoptotic cells. Genetic and pharmacological inhibition of SUV39H1/H2, the methyltransferases that deposit H3K9me3, resulted in upregulation of the pro-efferocytotic RBPJ-dependent genes and restoration of efferocytosis in *RBPJ*^-/-^ macrophages. These findings highlight a specific function of RBPJ in inflammation resolution and tissue repair governed by efferocytosis by promoting H3K9 trimethylation.

## Results

### RBPJ deficiency impairs efferocytosis and augments necrotic core areas in atherosclerotic plaques

We found increased *Rbpj* expression during inflammation driven by oxidized low-density lipoprotein (oxLDL) in bone-marrow-derived macrophages (BMDM) (Figure 1A). Consistently, we also detected altered expression of *Rbpj* and its downstream genes in aortic macrophages isolated from the mice fed with an atherogenic diet compared to those sorted from the mice fed with a regular diet (Figure 1B-C). To assess if RBPJ expression increases *in vivo* in an inflammatory condition, we measured the expression of this transcription factor in atherosclerotic plaque macrophages. We observed that macrophages residing in atherosclerotic plaques of mice (Figure 1D) and humans (Figure 1E) had heightened expression of RBPJ compared to the ones in normal arteries. To examine whether macrophage RBPJ plays a key role in inflammation propagation, we generated *Ldlr*^-/-^ *LyzM^cre/+^ Rbpj^fl/fl^* mice by bone marrow transplantation, and we validated the deletion of RBPJ in the BMDM of these mice (Supplemental Figure 1A). We observed that *Rbpj* deficiency did not alter apoptosis of macrophages (Supplemental Figure 1B). The frequencies (Supplemental Figure 1C) and numbers (Supplemental Figure 1D) of different leukocyte subsets in the aortas of these mice were similar to those in *Ldlr*^-/-^ *LyzM^+/+^ Rbpj^fl/fl^* littermate control mice. Interestingly, the frequency of Ly-6C^high^ monocytes was higher in the aortas of myeloid *Rbpj*-deficient mice. Additionally, we found no differences in the expression of the genes encoding inflammatory cytokines and chemokines, which exacerbate atherosclerosis, in the aortic arches of these mice (Supplemental Figure 1E). Similarly, these mice also had unaltered levels of serum cholesterol (Supplemental Figure 1F). *Rbpj* deficiency did not change the plaque area and fibrous cap thickness (Supplemental Figure 1G). However, atherosclerotic plaques in *Ldlr*^-/-^ *LyzM^cre/+^ Rbpj^fl/fl^* mice had larger necrotic core areas compared to the control mice (Figure 1F). We also validated this finding in *ApoE^-/-^* mice treated with siControl and si*Rbpj* incorporated in lipidoid nanoparticles, which are preferentially engulfed by macrophages (Supplemental Figure 1H) as we previously showed (37). Like genetic deficiency of *Rbpj* in myeloid cells, we found an increase in necrotic core area in si*Rbpj-*treated mice compared to siControl-treated mice while there was no change in fibrous cap thickness and plaque area (Figure 1G and Supplemental Figure 1I). Necrotic core areas in atherosclerotic plaques are formed partly due to defective clearance of apoptotic cells by foam cells, a process called efferocytosis (38). To assess defective efferocytosis *in vivo*, we performed a TUNEL assay in atherosclerotic plaques of myeloid *Rbpj*-deficient mice and observed diminished ratios of macrophage-associated non-nuclear TUNEL staining to macrophage-free TUNEL staining (Figure 1H). In line with this, we also observed that plaques from myeloid *Rbpj*-deficient mice harbored lower number of macrophages that engulfed caspase 3^+^ apoptotic particles (Figure 1I and Supplemental Figure 1J)(39). These data suggest that *Rbpj*- deficient macrophages have impaired ability to clear apoptotic cells.

**Figure 1:**
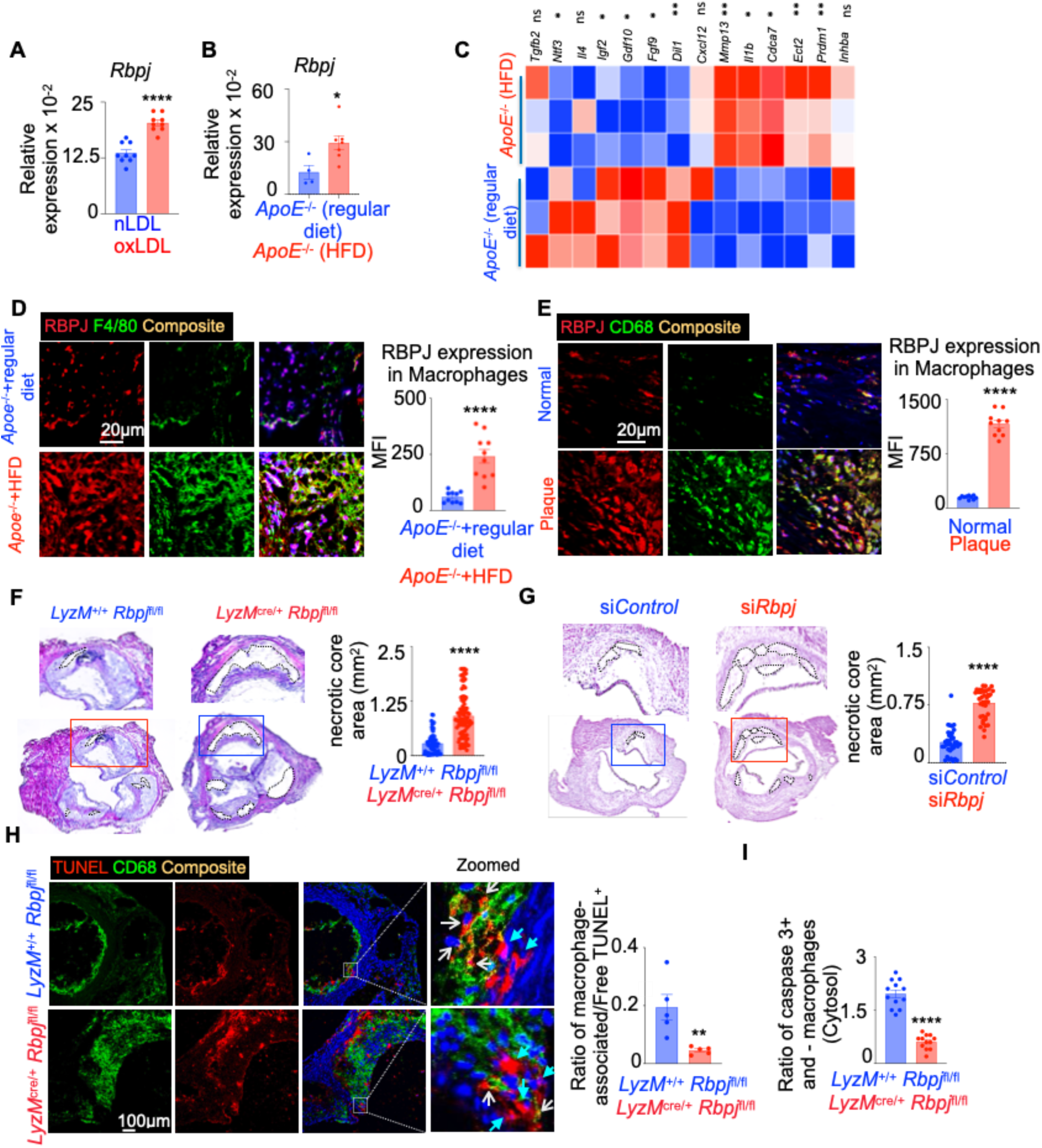
RBPJ is crucial for macrophage efferocytosis. **A.** qPCR quantification of *Rbpj* in native LDL and ox-LDL-treated BMDM. (n=7-9/group). **B-C.** *ApoE*^-/-^ mice were fed with either a regular or high fat diet (HFD) for 16 weeks. Aortas were harvested, and plaque-associated macrophages were sorted by FACS. qPCR was performed to quantify *Rbpj* (**B**) and its downstream genes represented as a heat map showing three replicates (**C**) (n=5-7/group). **D**. Representative confocal microscopy images of aortic root sections of *Apoe*^-/-^ mice fed with a regular or high fat diet and quantification of RBPJ expression in plaque macrophages. (n=5/group). **E.** Representative confocal microscopy images of normal and plaque-containing carotid arteries of humans and quantification of RBPJ expression in plaque macrophages. (n=5 group). **F.** Representative H&E staining images of atherosclerotic plaques in aortic root sections of *Ldlr*^-/-^ *LyzM^+/+^ Rbpj^fl/fl^* and *Ldlr*^-/-^ *LyzM^cre/+^ Rbpj^fl/fl^* mice and quantification of necrotic core areas. (n=4-5/group). **G.** Representative H&E staining images of the aortic roots of si*Control* and si*RBPJ*-treated *ApoE^-/-^* mice fed with an HFD (n=6/group). **H**. To assess *in situ* efferocytosis, aortic root sections of *Ldlr*^- /-^ *LyzM*^+/+^ *Rbpj*^fl/fl^ and *Ldlr*^-/-^ *LyzM^cre^*^/+^ *Rbpj*^fl/fl^ mice were stained with anti-CD68 antibody (green) to label macrophages and with TUNEL stain (red) to detect apoptotic cells. The ratios of macrophage-associated non-nuclear TUNEL staining (white arrows) to macrophage-free TUNEL staining (blue arrows) were calculated to determine efferocytosis within the lesions. (n=5/group). **I.** Quantification of caspase 3 mean fluorescence intensity (MFI) in aortic root plaque macrophages of *Ldlr*^-/-^ *LyzM^+/+^ Rbpj^fl/fl^* and *Ldlr*^-/-^ *LyzM^cre/+^ Rbpj^fl/fl^* mice. (n=4-5/group). The data are expressed as mean ± SEM and from 3–5 independent experiments. \*\**p* < 0.05, \*\**p* < 0.01, \*\*\*\**p* < 0.0001.

### RBPJ deletion dampens macrophage efferocytosis

To investigate if RBPJ deficiency in macrophage compromises efferocytosis, BMDM from *LyzM^+/+^ Rbpj^fl/fl^* and *LyzM^cre/+^ Rbpj^fl/fl^* mice were cultured in presence of apoptotic Jurkat cells, fluorescently labeled with PKH67 dye, and evaluated for the uptake of apoptotic particles (Figure 2A). *Rbpj*-deficient macrophages exhibited lower apoptotic cell engulfment than control macrophages at 37° C, the temperature amicable for efferocytosis (Figures 2B-C and Supplemental Figure 2A). The experiment was then carried out at 4° C, which promotes apoptotic cell attachment to macrophages but inhibits their engulfment(40). BMDM from *LyzM^cre/+^ Rbpj^fl/fl^* mice exhibited lower apoptotic cell attachment compared to control mice (Figure 2B), indicating that defective macrophage efferocytosis of Rbpj-deficient mice was partially due to diminished binding of apoptotic cells. To confirm impaired binding and engulfment of apoptotic cells in absence of macrophage RBPJ, we also did a one-hour time lapse imaging of efferocytosis by WT and *Rbpj* KO BMDM. We found a slow engulfment of apoptotic cells by *Rbpj* KO BMDM (Figure 2D and Supplemental Videos 1 and 2). To confirm these findings, we silenced *Rbpj* in BMDM and cultured them in the presence of apoptotic Jurkat cells labelled with a pHrodo dye. We found 4-fold decrease in apoptotic cell engulfment and 2-fold decrease in binding after *Rbpj* silencing (Supplemental Figure 2B-C). Furthermore, *Rbpj* deficiency decreased the genes facilitating efferocytosis (Figure 2E). Of note, macrophages upregulated *Rbpj* expression after they were exposed to apoptotic cells (Supplemental Figure 2D). RBPJ is the principal effector of the Notch signaling pathway, and the cleavage of the Notch receptor by γ-secretase to form Notch intracellular domain (NICD) is crucial for the downstream transcriptional activation(41, 42). We used DAPT, a well-known inhibitor of γ-secretase(43), to examine whether RBPJ mediated apoptotic cell clearance is dependent on the Notch signaling. A significant reduction in apoptotic Jurkat cell engulfment was observed in BMDM treated with DAPT (Figure 2F). Conversely, the Notch activator Yhhu-3792(44) further elevated apoptotic cell engulfment in *Rbpj*^+/+^ macrophages suggesting the merit of the Notch signaling in efferocytosis (Figure 2G). To understand if the impairment of efferocytosis due to Notch inhibition can be reversed by RBPJ signaling, we overexpressed *RBPJ* in THP-1 macrophages after inhibiting the Notch signaling with DAPT (Supplemental Figure 2E). RBPJ overexpression led to a significant increase in efferocytosis in the DAPT-treated cells (Figure 2H-I and Supplemental Figure 2F)

**Figure 2:**
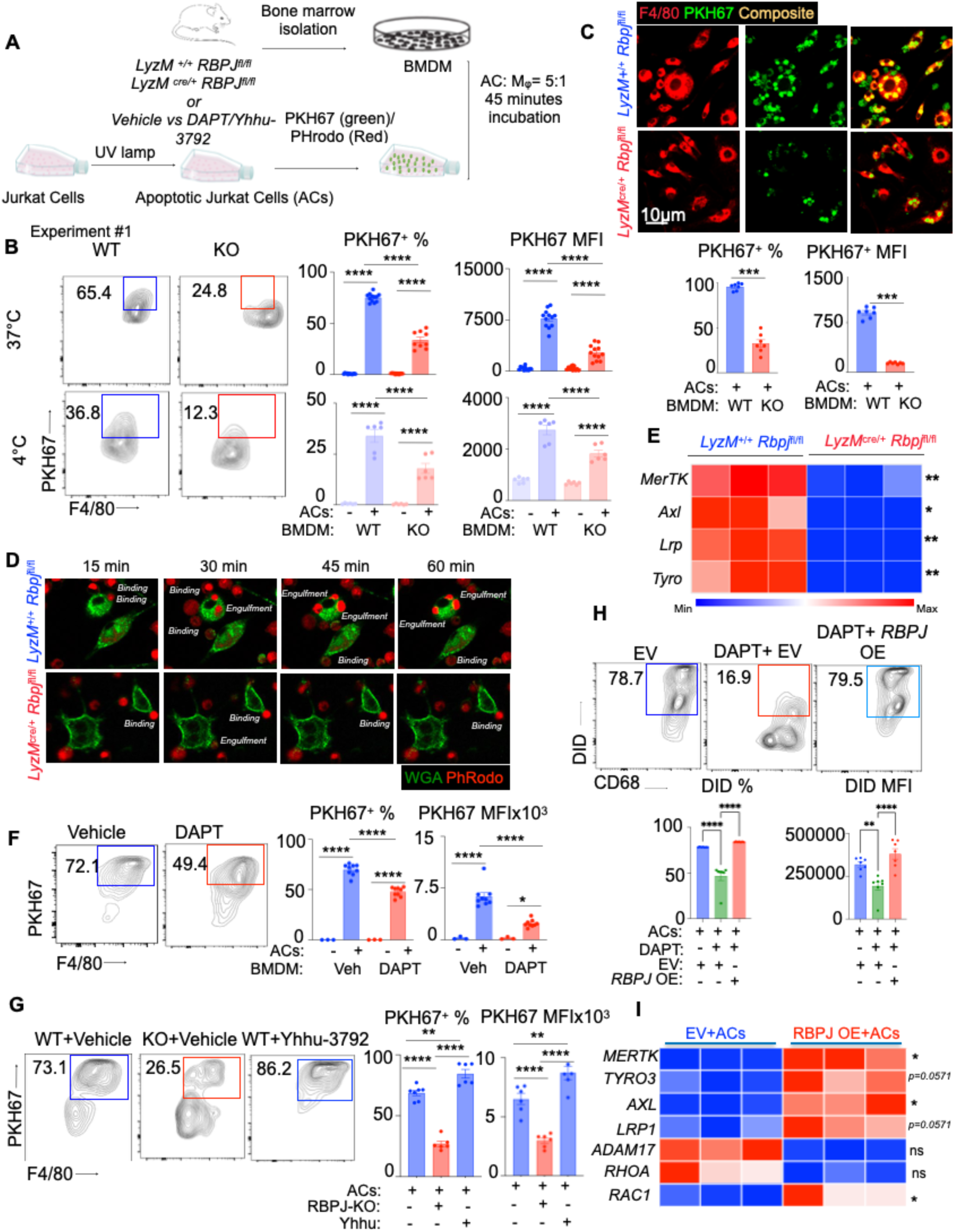
BMDM-mediated efferocytosis depends upon the Notch-RBPJ signaling cascade. **A.** Schematic diagram of the experiments understanding the role of the Notch-RBPJ signaling pathway in mice bone marrow derived macrophage (BMDM)-mediated efferocytosis. **B.** Flow cytometry plots showing the uptake of PKH67^+^ apoptotic Jurkat cells by WT and *Rbpj* KO BMDM and the quantification of the frequency of PKH67^+^ BMDM and PKH67 MFI at 37^0^ C and 4^0^ C. (n=6 /group). **C.** Representative confocal microscopy images showing engulfment of PKH67-labeled apoptotic Jurkat cells by BMDM of *Rbpj*^+/+^ and *Rbpj*^-/-^ mice and the quantification of % of PKH67^+^ BMDM and PKH67 mean fluorescence intensities (MFI). (n=7-8 /group). **D.** Time lapse images showing binding and engulfment of pHRodo labelled ACs by WT and *Rbpj* KO BMDM (n=5/group)**. E.** Heatmap showing three qPCR replicates quantifying efferocytosis-related genes in *Rbpj*⁺/⁺ and *Rbpj*⁻/⁻ BMDMs in the presence of apoptotic Jurkat cells. (n=5-10/group). **F.** Flow cytometry plots showing the uptake of PKH67^+^ apoptotic Jurkat cells and the quantification of the frequency of PKH67^+^ BMDM and PKH67 MFI in BMDM treated with either vehicle or DAPT (Notch inhibitor) at 37^0^ C. (n=3-9/group). **G.** Flow cytometry plots showing the uptake of PKH67^+^ apoptotic Jurkat cells and the quantification of the frequency of PKH67^+^ BMDM and PKH67 MFI in *Rbpj*^+/+^ and *Rbpj*^-/-^ BMDM treated with either vehicle or Yhhu-3792 (Notch activator) at 37^0^ C. (n=5-7/group). **H-I**. THP-1 macrophages were treated with DAPT (Notch inhibitor) and infected with either empty vector (EV) or *RBPJ* lentivirus (*RBPJ* OE). The cells were co-cultured with DiD-labeled apoptotic Jurkat cells (ACs). **H.** The representative flow cytometry plots and the bar graphs showing efferocytosis quantification (n=7/group). **I.** The heatmap shows three replicates of qPCR quantification of the efferocytotic genes in control and RBPJ-overexpressing THP-1 macrophages. (n=7/group). The data are expressed as mean ± SEM and from 3 independent experiments. \**p* < 0.05, \*\**p* < 0.01, \*\*\**p* < 0.001, \*\*\*\**p* < 0.0001.

### *Rbpj* deletion reduces efferocytic capacity of peritoneal macrophages in acute peritonitis and alveolar macrophages in acute lung injury

We used zymosan-induced peritonitis, a well-known model of acute inflammation and resolution, to test the hypothesis that RBPJ increases efferocytosis during inflammation resolution. Low-dose zymosan elicits a neutrophil-mediated inflammatory response, apoptosis of neutrophils, and clearance of dead cells by peritoneal macrophages(45). To assess *in vivo* efferocytic potential of peritoneal macrophages, we injected *LyzM^+/+^ Rbpj^fl/fl^* and *LyzM^cre/+^ Rbpj^fl/fl^* mice with fluorescently labeled apoptotic neutrophils in the peritoneal cavity (Figure 3A). *Rbpj*-deficient mice exhibited significantly reduced uptake of apoptotic neutrophils by peritoneal macrophage. (Figures 3B and 3C). The peritoneal cavity harbors two distinct macrophage subsets with phagocytic ability: large (CD11b^high^ F4/80^high^) and small (CD11b^low^ F4/80^low^) peritoneal macrophages(46). We observed that the ability of engulfing apoptotic neutrophils of both macrophage subsets was curtailed due to *Rbpj* deficiency (Figure 3D). Furthermore, the peritoneal exudate fluid in *LyzM^cre/+^ Rbpj^fl/fl^* mice contained higher amount of IL-1β (Figure 3E). In line with this, flow cytometric analysis of *Rbpj*-deficient BMDMs exposed to apoptotic cells revealed increased TNF-α and decreased IL-10 expression (Supplemental Figure 2G). Also, when RBPJ was silenced in BMDM, they exhibited increased expression of inflammatory gene *Il1b* and *Il6* while anti-inflammatory genes like *Il10* and *Tgfb* were downregulated (Supplemental Figure 2H). These findings indicate that in the context of impaired efferocytosis, *Rbpj* deficiency skews macrophages toward a pro-inflammatory phenotype. Neutrophil apoptosis is a hallmark of diseases such as acute peritonitis and acute lung injury (ALI) (47, 48). Similar to Jurkat cells, the efferocytosis of neutrophils was also impaired in Rbpj deficiency (Supplemental Figure 3A-B).

**Figure 3:**
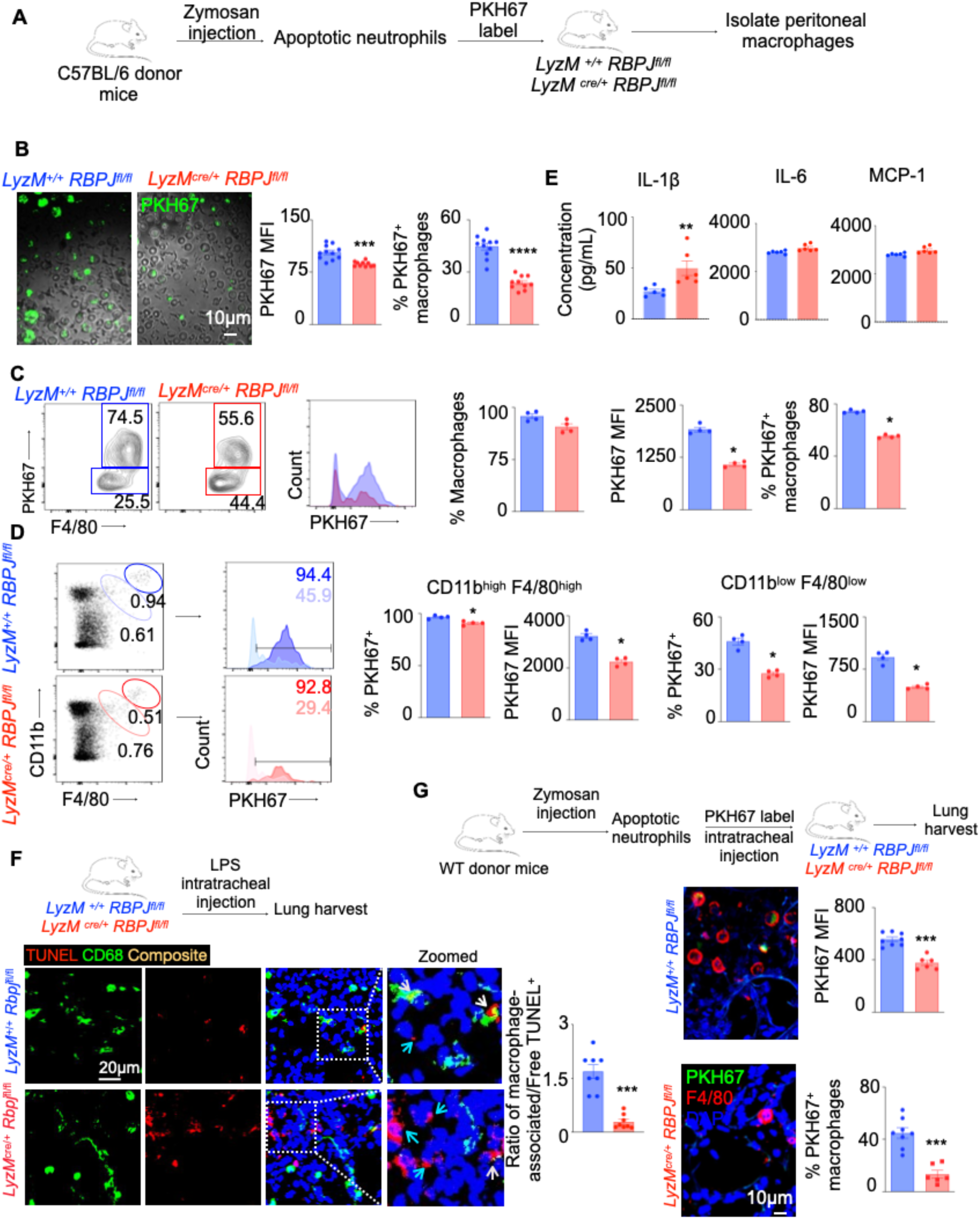
RBPJ promotes the efferocytosis capacity of peritoneal macrophages and alveolar macrophages *in vivo.* **A.** Schema showing the experimental design to test the effect of *Rbpj* deficiency in peritoneal macrophages on efferocytosis. **B.** Representative microscopy images of PKH67-labeled apoptotic neutrophil engulfment by peritoneal macrophages in *LyzM^+/+^ Rbpj^fl/fl^* and *LyzM^cre/+^ Rbpj^fl/fl^* mice and quantification of PKH67 MFI and % of PKH67^+^ macrophages. (n=10-11 /group). **C.** Flow cytometry plots and quantification of PKH67^+^ apoptotic neutrophil uptake by peritoneal macrophages in *LyzM^+/+^ Rbpj^fl/fl^* and *LyzM^cre/+^ Rbpj^fl/fl^* mice. (n=4 /group). **D.** Flow cytometry plots and quantification of PKH67^+^ apoptotic neutrophil uptake by CD11b^high^ F4/80^high^ and CD11b^low^ F4/80^low^ peritoneal macrophages in *LyzM^+/+^ Rbpj^fl/fl^* and *LyzM^cre/+^ Rbpj^fl/fl^*mice. (n=4/group). **E.** Quantification of the cytokines by ELISA in peritoneal exudates. (n=6/group). **F.** Experimental design to assess apoptotic cells in the lungs of *LyzM^+/+^ Rbpj^fl/fl^* and *LyzM^cre/+^ Rbpj^fl/fl^* mice. Representative confocal images showing CD68 (green) and TUNEL (red) staining and bar graphs quantifying the ratios of macrophage-associated non-nuclear TUNEL staining (white arrows) to macrophage-free TUNEL staining (blue arrows) to measure efferocytosis by lung macrophages. (n=8/group). **G.** Experimental design to discern the effects of RBPJ on alveolar macrophage-mediated efferocytosis. The data are expressed as mean ± SEM and obtained from 3 independent experiments. \**p* < 0.05, \*\**p* < 0.01, \*\*\**p* < 0.001, \*\*\*\**p* < 0.0001.

ALI is accompanied by widespread apoptosis of alveolar epithelial cells(49). Additionally, evidence from murine models and clinical studies suggests that effective efferocytosis after ALI is essential for lung healing.(50). To examine whether RBPJ stimulates macrophage efferocytosis following ALI, we exposed *LyzM^+/+^ Rbpj^fl/fl^* and *LyzM^cre/+^ Rbpj^fl/fl^* mice to LPS-induced ALI (Figure 3F). Immunofluorescent imaging confirmed efficient reduction of RBPJ-expressing lung macrophages in *LyzM^cre/+^ Rbpj^fl/fl^* mice (Supplemental Figure 3C). Analysis of *in situ* efferocytosis employing TUNEL staining in lung sections of these mice revealed diminished ratios of non-nuclear macrophage-associated to macrophage-free TUNEL staining in lungs from *Rbpj*-deficient mice, indicating impaired efferocytosis in the absence of myeloid *Rbpj* (Figure 3F). We did not find any alterations in nuclear TUNEL staining in these mice suggesting that macrophage cell death is not increased in *Rbpj*-deficient mice (Supplemental Figure 3D). Moreover, mice deficient of macrophage *Rbpj* had increased frequency of caspase 3^+^ cells (outside of macrophages) in the lungs compared to the littermate control mice, suggesting that apoptotic bodies not being efficiently cleared by *Rbpj*-deficient macrophages (Supplemental Figure 3E). To further investigate whether RBPJ aids in efferocytosis by alveolar macrophages, we intratracheally injected fluorescent labeled apoptotic neutrophils (Figure 3G). The engulfment of the injected apoptotic neutrophils by alveolar macrophages was significantly reduced in absence of macrophage *Rbpj* (Figure 3G). As a whole, by using these different mouse models of efferocytosis, we found that *Rbpj* deletion decreases efferocytic capacity of macrophages *in vivo*.

### *Rbpj* deficiency alters the transcriptional profile of macrophages during efferocytosis

To uncover the genes implicated in RBPJ-mediated efferocytosis, we conducted RNA sequencing in peritoneal macrophages of *LyzM^+/+^ Rbpj^fl/fl^* and *LyzM^cre/+^ Rbpj^fl/fl^* mice intraperitoneally injected with apoptotic neutrophils (Figure 4A). For control, we intraperitoneally injected *LyzM^+/+^ Rbpj^fl/fl^*mice with PBS. Our RNA seq data confirmed a four-fold decrease in *Rbpj* expression in peritoneal macrophages obtained from *LysM*^Cre/+^ *Rbpj*^fl/fl^ mice compared to those derived from *LysM*^+/+^ *Rbpj*^fl/fl^ mice (Supplemental Figure 3F). We also observed *Mertk* which facilitates efferocytosis to be upregulated in WT+ACs vs WT+PBS (Supplemental Figure 3G). Whole transcriptome analysis revealed that efferocytosis altered the transcriptomic profile of peritoneal macrophages as previously reported(5, 40, 45, 51, 52) (Figures 4B-C). Presence of apoptotic cells changed the expression of 1257 genes in peritoneal macrophages. For example, StAR-related lipid transfer domain protein 13 (STARD13) can selectively activate RhoA and Cdc42 and inhibit cell growth by suppressing actin stress fiber assembly(53). EF-hand calcium-binding domain-containing protein 5 (EFCAB5) regulates brain aging as well as calcium signal transduction, synapse generation, and cell survival(54). *Rbpj* deletion changed the transcriptomic profile of peritoneal macrophages during efferocytosis as shown in the PCA analysis, volcano plot, and heatmap (Figures 4B-4D). We identified 261 genes that were differentially expressed in *Rbpj*-deficient macrophages compared to wildtype macrophages in presence of apoptotic cells. Among these genes, 49 were also differentially expressed in wildtype peritoneal macrophages in response to apoptotic cells (Figure 4E). When we compared the expression of these 49 genes in the three treatment groups, we observed that they formed four distinct clusters (Figure 4F). The Cluster 1 genes (Figure 4G) were of particular interest because the expression of these genes was heightened in wildtype macrophages in presence of apoptotic cells while their expression was dampened in absence of *Rbpj*. From these data, we hypothesized that the Cluster 1 genes, hereon referred to as RBPJ-dependent genes, regulate RBPJ-mediated efferocytosis.

**Figure 4:**
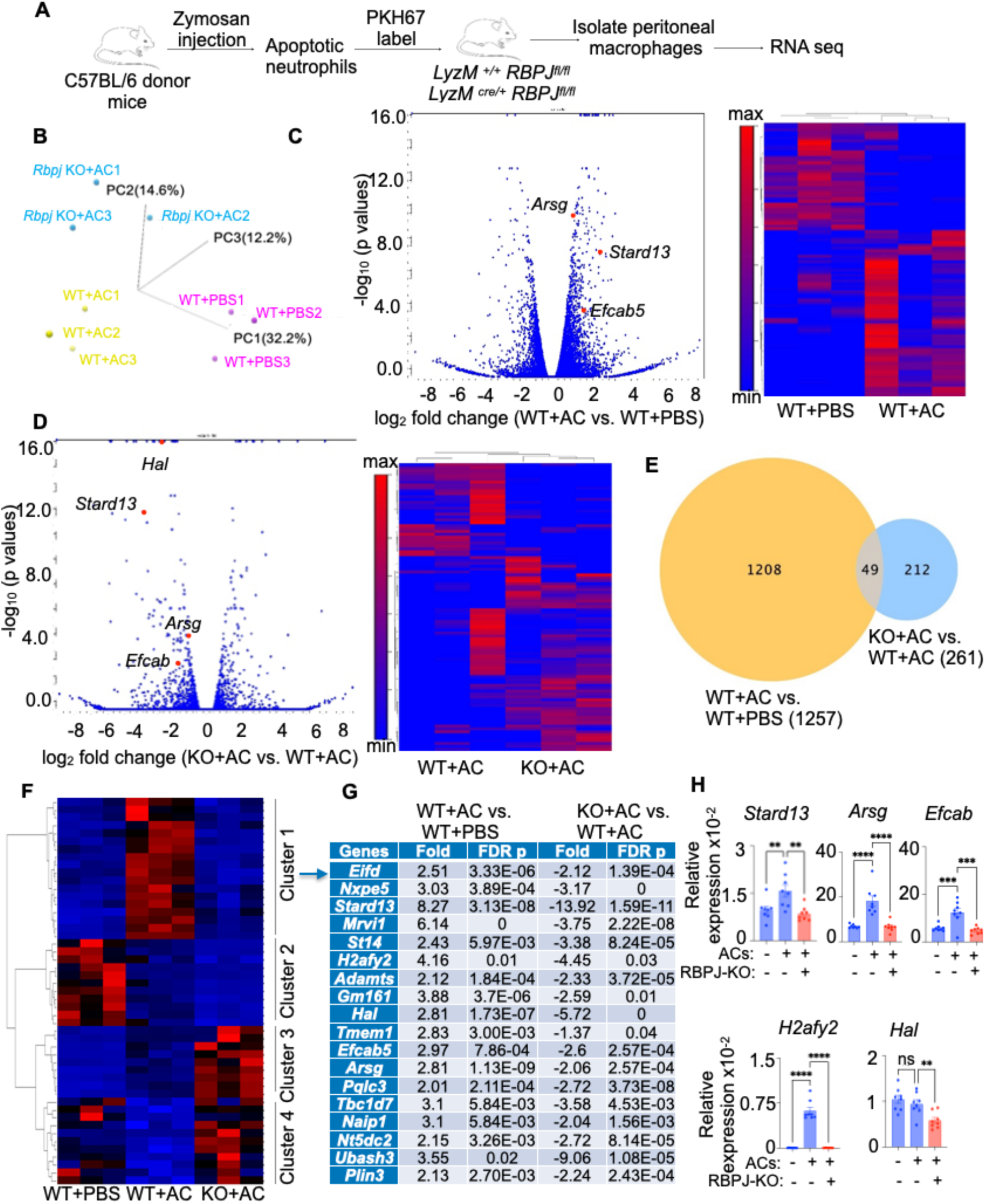
*Rbpj* deficiency alters efferocytosis-associated transcriptomic profile in macrophages. **A.** Bulk RNA sequencing on *Rbpj*^+/+^ (WT) and *Rbpj*^-/-^ (KO) peritoneal macrophages exposed or non-exposed to ACs. **B-D.** Principal component analysis (PCA) of the transcriptomic profiles (**B**) and the volcano plot and heatmap showing differentially expressed genes (**C-D**) of WT peritoneal macrophages exposed or non-exposed to ACs and *Rbpj* KO peritoneal macrophages exposed to ACs. (n=3 /group). **E.** Venn diagram representing 49 overlapping genes between 1257 differentially expressed genes in WT+ACs vs. WT+PBS and 261 differentially expressed genes in KO+ACs vs. WT+ACs. (n=3 /group). **F.** Classification of the 49 overlapping genes in various clusters. (n=3 /group). **G.** Fold changes and FDR p values of the Cluster 1 genes comparing 1. *RBPJ*^+/+^ peritoneal macrophages exposed to ACs and those not exposed to ACs and 2. AC-exposed *Rbpj*^-/-^ vs. *Rbpj*^+/+^ peritoneal macrophages. (n=3 /group). **H.** qPCR quantification of the Cluster 1 genes or RBPJ-dependent genes in *Rbpj*^+/+^ and *Rbpj*^-/-^ BMDM cultured with or without apoptotic Jurkat cells (n=5-10 /group). The data from an RNA sequencing experiment are shown. qPCR data are expressed as mean ± SEM \*\**p* < 0.01, \*\*\**p* < 0.001, \*\*\*\**p* < 0.0001.

### *Rbpj* deficiency increases H3K9me3 in the promoter regions of the RBPJ-dependent genes

Since RBPJ is known to regulate gene expression by directly binding to promoter regions(19, 22), we performed *in silico* analysis to discern whether RBPJ has binding motifs in the promoter regions of the RBPJ-dependent genes. We observed that the *Stard13, Arsg*, *Efcab5*, *H2afy2*, and *Hal* promoters include RBPJ binding sequences (Supplemental Figure 4A). Quantitative PCR confirmed upregulation of most of these genes during efferocytosis while their expression was downregulated in absence of *Rbpj* (Figure 4H). Using RBPJ ChIP followed by qPCR (ChIP-qPCR), we confirmed the interactions between this transcription factor and the *Stard13, Arsg*, *Efcab5*, *H2afy2*, and *Hal* promoters in the presence (Supplemental Figure 4B) or absence (Figure 5A) of apoptotic cells. Epigenetic modifications license tissue specific functions of resident macrophages(51, 52, 55). To understand if efferocytosis depends on epigenetic reprogramming of macrophages, we have evaluated the expression of H4AC, a gene activation marker, and H3K9me3, a gene repression marker in presence or absence of apoptotic cells. In line with increased expression of the RBPJ-dependent genes in presence of apoptotic cells, H3K9me3 in the promoter regions of these genes in macrophages cultured with apoptotic Jurkat cells decreased (Figure 5B) while H4AC elevated in the *Stard13* and *H2afy2* promoters (Supplemental Figure 4C), indicating activation of *Stard13, Arsg*, *Efcab5*, *H2afy2*, and *Hal* during efferocytosis. Additionally, we observed augmented H3K9me3 in the promoter regions of these genes in the absence of *Rbpj* in macrophages (Figure 5B) while H4AC was unchanged (Supplemental Figure 4C). Since the ChIP-qPCR experiments demonstrated reduced H3K9me3 expression in the promoter regions of the RBPJ-dependent genes during efferocytosis, we investigated global genomic H3K9me3 deposition in promoter regions using Cleavage Under Targets and Release Using Nuclease (CUT&RUN) analysis during macrophage efferocytosis in the presence or absence of RBPJ. We observed H3K9me3 marks at different regions of genes (Supplemental Figure 4D). Remarkably, the H3K9me3 peaks were higher in the absence of RBPJ in macrophages undergoing efferocytosis (Figure 5C, Supplemental Figures 4E and 4F). Next, we evaluated the H3K9me3 mark in the RBPJ-dependent genes, and we found that this methylation was reduced in the presence of RBPJ during efferocytosis (WT+ACs compared to WT+PBS) while *Rbpj* deficiency increased H3K9me3 marks in the RBPJ-dependent genes (KO+ACs compared to WT+ACs) (Figure 5D). When we compared the genes having the H3K9me3 mark in resting *Rbpj*^+/+^ macrophages, and *Rbpj*^+/+^ and *Rbpj*^-/-^ macrophages mediating efferocytosis, 245 unique genes were identified in *Rbpj*^-/-^ macrophages undergoing efferocytosis (Figure 5E). These genes are enriched in cellular functions crucial to efferocytosis, such as cell interaction, cell adhesion, cell projection, and cell junction assembly (Figure 5F). Similarly, we discerned the genes that are common between A) genes with the H3K9me3 mark in *Rbpj*^-/-^ macrophages compared to *Rbpj*^+/+^ macrophages undergoing efferocytosis and B) upregulated genes in *Rbpj*^+/+^ macrophages undergoing efferocytosis (Supplemental Figure 4G). These common genes are reported to be involved in negative regulation of DNA binding, protein kinase B signaling, protein import, contact inhibition etc. (Supplemental Figure 4H). We also performed CUT&RUN sequencing for H3K9me3 in WT and *Rbpj^-/-^*BMDM at baseline (PBS) (Supplemental Figure 5). Supplemental Figure 5A shows a genome-wide distribution of H3K9me3. We did not observe any significant differences in H3K9me3 enrichment intensities in various regions of the genome. These results suggest that the absence of RBPJ does not alter baseline H3K9me3 levels, but instead contributes to increased H3K9me3 expression specifically in the presence of apoptotic cells (Supplemental Figure 5B-E). We observed a significant increase in the frequency of H3K9me3-expressing macrophages in atherosclerotic plaques of mice lacking *Rbpj* in myeloid cells (Figure 5G). To experimentally validate the role of H3K9me3 in RBPJ-driven efferocytosis, we silenced histone methyltransferases *Suv39h1/h2*, responsible for this epigenetic alteration, using siRNAs against both *Suv39h1* and *Suv39h2* methyl transferases in *Rbpj*^+/+^ and *Rbpj*^-/-^ BMDM undergoing efferocytosis (Supplemental Figure 6A and 6B). Interestingly, we observed no change in efferocytosis in *Rbpj*^-/-^ BMDM treated with either *Suv39h1* or *Suv39h2* siRNAs as compared to siControl-treated *Rbpj*^-/-^ BMDM (Figure 5H and Supplemental Figure 6C). However, silencing both histone methyltransferases reversed the lower efferocytosis in absence of *Rbpj*. We observed similar reversal of apoptotic cell debris uptake by macrophages after the treatment with chaeotocin(56), which inhibits both histone methyltransferases (Supplemental Figure 6D). Additionally, *Suv39h1*/*Suv39h2* siRNAs (Figure 5I) and chaeotocin (Supplemental Figure 6E) treatment restrained downregulation of the RBPJ-dependent genes in the absence of *Rbpj*. Finally, among the histone marks we checked, only H3K9me3 expression was augmented in *Rbpj*^-/-^ macrophages (Supplemental Figure 6F). In cluster, these data suggest that RBPJ dampens H3K9me3, increasing the expression of the RBPJ-dependent genes, to facilitate efferocytosis. We also assessed pro- and anti-inflammatory genes (*Il1b, Tnfa, IL-6, Il10*, and *Tgfb*) after chaotocin and *Suv39h1*/*Suv39h2* siRNAs treatment in BMDM in the presence of apoptotic Jurkat cells. We observed that pro-inflammatory genes, such as *Il6* and *Il1b,* were significantly downregulated in *Rbpj* KO BMDM treated with a combination of both the siRNAs compared to *Rbpj* KO BMDM alone (Supplemental Figure 6G). A similar trend was observed following chaetocin treatment of *Rbpj* KO BMDM (Supplemental Figure 6H). Notably, while the pro-inflammatory gene *Il6* was downregulated, anti-inflammatory genes such as *Il10* and *Tgfb* were upregulated. Altogether, these data suggest that loss of RBPJ leads to increased H3K9me3 deposition in the promoters of the RBPJ-dependent genes, resulting in their reduced expression.

**Figure 5:**
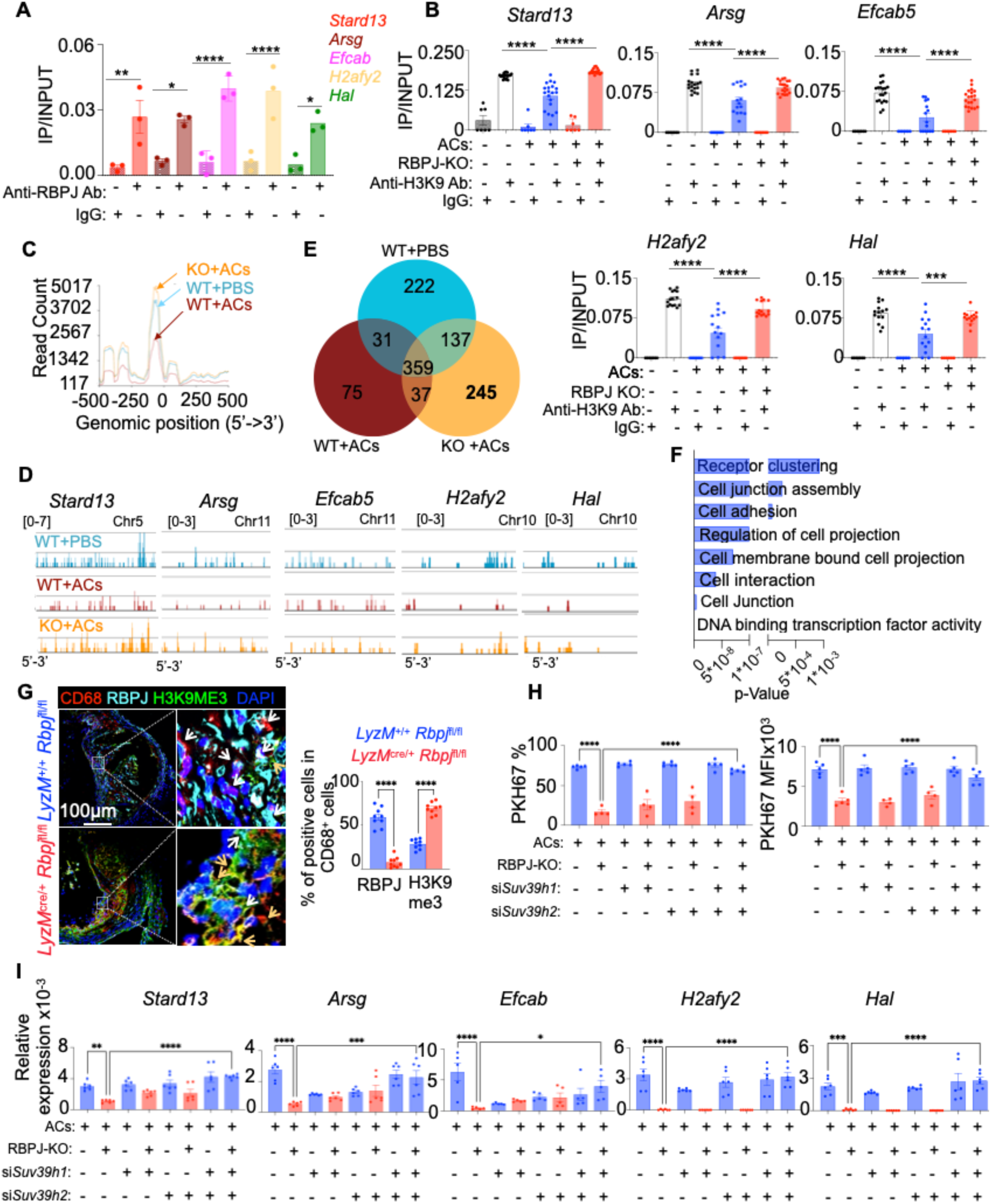
RBPJ deficiency facilitates H3K9me3 enrichment in the RBPJ-dependent genes during efferocytosis. **A.** ChIP-qPCR quantification of RBPJ occupancy on the *Stard13, Arsg, Efcab5, H2afy2*, and *Hal* promoters in *RBPJ*^+/+^ BMDM cultured in absence of apoptotic cells (n=3/group). One of two representative experiments is shown. **B.** ChIP-qPCR quantification of H3K9me3 mark in the promotor regions of *Stard13, Arsg, Efcab5, H2afy2*, and *Hal* in *RBPJ*^+/+^ and *RBPJ*^-/-^ BMDM in the presence or absence of apoptotic Jurkat cells. (n=5-10 /group). **C.** Intensity plot of CUT&RUN analysis to show H3K9me3 enrichment around +/-500 bp of the transcription start sites. (n=3/group). **D.** CUT&RUN analysis showing H3K9me3 peak distribution of the RBPJ-dependent genes: *Stard13, Arsg, Efcab5, H2afy2,* and *Hal*. (n=3/group). **E.** Venn diagram depicting 245 unique H3K9me3-enriched regions in *RBPJ*^-/-^ BMDM exposed to apoptotic cells (KO+ACs) when compared with *RBPJ*^+/+^ BMDM exposed (WT+ACs) or non-exposed (WT+PBS) to apoptotic cells. (n=3 /group). **F.** Gene set enrichment analysis of the 245 unique genes obtained from the Venn diagram analysis. (n=3/group). **G.** Representative confocal images of aortic root sections of *Ldlr*^-/-^ *LyzM^+/+^ Rbpj^fl/fl^* and *Ldlr*^-/-^ *LyzM^cre/+^ Rbpj^fl/fl^* mice showing RBPJ and H3K9me3 expression in plaque macrophages. The frequencies of RBPJ^+^ (white arrow) and H3K9me3^+^ (yellow arrow) macrophages are shown. (n=5/group). **H.** Flow cytometry-assisted quantification showing PKH67^+^ apoptotic Jurkat cell uptake by si*control*, si*Suv39h1,* si*Suv39h2*, and si*Suv39h1/* si*Suv39h2-*transfected BMDM of *Rbpj*^+/+^ and *Rbpj*^-/-^ mice. (n=4-5/group). **I.** qPCR quantification of the RBPJ-dependent genes in the BMDM obtained from H. (n=4-6/group). The CUT&RUN data are obtained from one experiment. The other data are obtained from 3 independent experiments. The data are expressed as mean ± SEM. \**p* < 0.05, \*\**p* < 0.01, \*\*\**p* < 0.001, \*\*\*\**p* < 0.0001.

**Figure 6:**
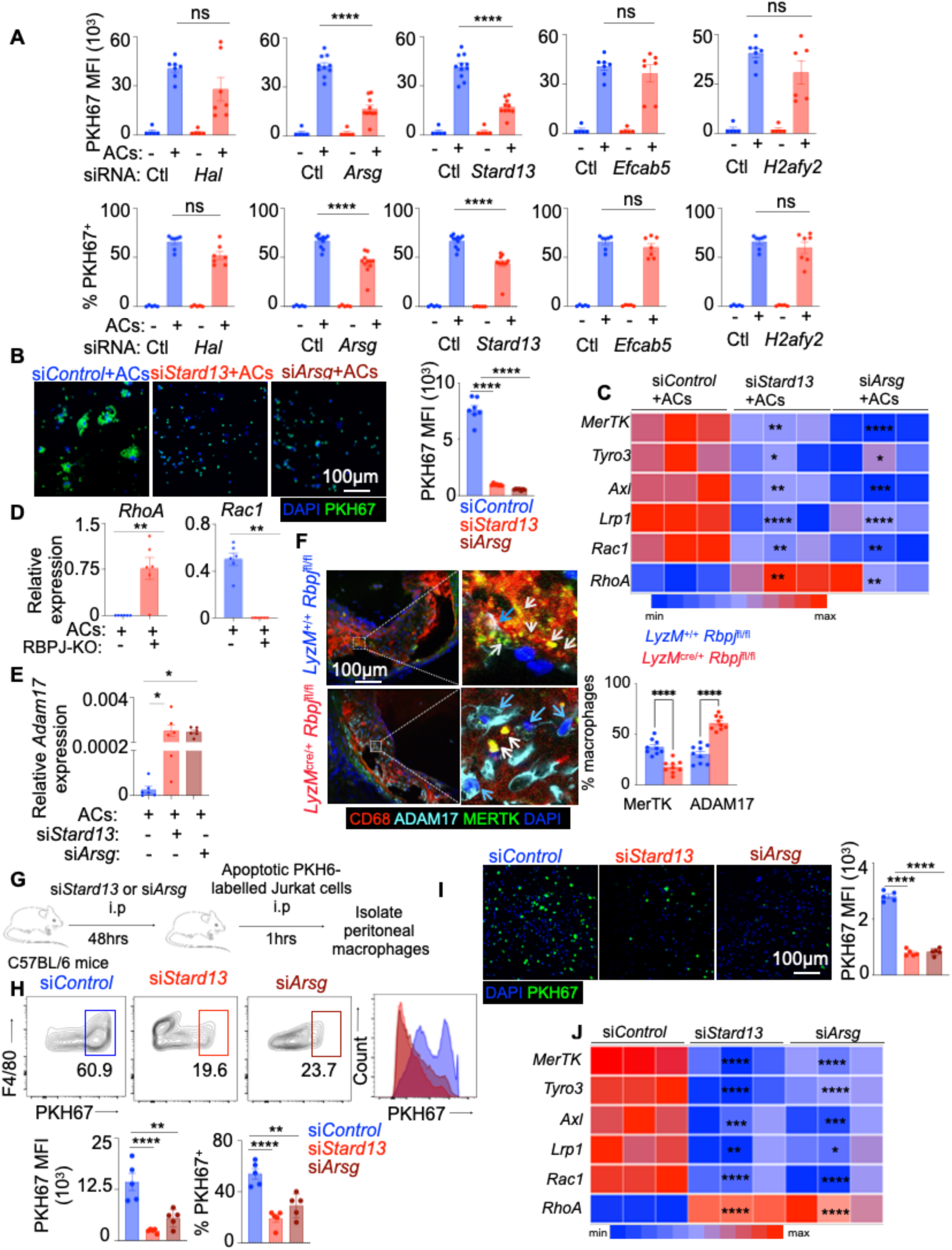
RBPJ-mediated efferocytosis is driven by Stard13 and Arsg. **A.** Quantification of efferocytosis by BMDM in the presence or absence of PKH67^+^ apoptotic Jurkat cells after si*Control*, si*Hal*, si*Arsg*, si*Stard13*, si*Efcab*, or si*H2afy2* treatment using flow cytometry. (n=5-7/group). **B.** Representative microscopy images and quantification of efferocytosis by BMDM in presence of PKH67^+^ apoptotic Jurkat cells after si*Control*, si*Arsg*, or si*Stard13* treatment. (n=5-7/group). **C.** Heatmap showing three qPCR replicates quantifying the efferocytosis-related genes *MerTk, Tyro3, Axl, Lrp1, Rac1,* and *RhoA* in BMDM in the presence of apoptotic Jurkat cells after si*Control*, si*Arsg*, or si*Stard13* treatment. (n=5-10 /group). **D.** qPCR quantification of *RhoA* and *Rac1* in the presence of apoptotic Jurkat cells in *Rbpj*^+/+^ and *Rbpj*^-/-^ BMDM. (n=5-10/group). **E.** qPCR analysis of *Adam17* expression in BMDM transfected with si*Control*, si*Stard13*, or si*Arsg* (n=5-6/group). **F.** Representative confocal images of aortic root sections of *Ldlr*^-/-^ *LyzM*^+/+^ *Rbpj*^fl/fl^ and *Ldlr*^-/-^ *LyzM^cre^*^/+^ *Rbpj*^fl/fl^ mice showing MERTK (white arrow) and ADAM17 (blue arrow) expression in CD68^+^ cells (macrophages). The frequencies of MERTK^+^ and ADAM17^+^ macrophages are shown in the bar graphs. (n=5/group). **G.** Experimental design to evaluate the impact of Arsg and Stard13 on *in vivo* efferocytosis by peritoneal macrophages**. H.** Representative flow cytometry plots and quantification of efferocytosis by peritoneal macrophages after si*Control*, si*Arsg*, or si*Stard13* treatment. (n=5/group). **I.** Representative microscopy images of PKH67-labeled apoptotic Jurkat cell engulfment by peritoneal macrophages treated with *control, Stard13,* and *Arsg* siRNA and quantification of engulfed cells represented as PKH67 MFI. (n=5/group). **J.** Assessment of the expression of the efferocytosis genes-*MerTk, Tyro3, Axl, Lrp1, Rac1,* and *RhoA* in peritoneal macrophages isolated from mice injected with si*Control,* si*Stard13*, or *siArsg*. (n=5/group). The data are expressed as mean ± SEM and obtained from 3–5 independent experiments. \**p* < 0.05, \*\**p* < 0.01, \*\*\**p* < 0.001, \*\*\*\**p* < 0.0001.

### Stard13 and Arsg promote efferocytosis

We observed that RBPJ regulates the expression of the RBPJ-dependent genes *Stard13, Arsg*, *Efcab5*, *H2afy2*, and *Hal* and the promoter regions of these genes contain RBPJ binding motifs. From these data we hypothesized that the RBPJ-dependent genes promote efferocytosis. To this end, we silenced *Stard13, Arsg*, *Efcab5*, *H2afy2*, and *Hal* in BMDM (Supplemental Figure 7A). Apoptotic Jurkat cell engulfment in BMDM was significantly reduced after *Stard13* and *Arsg* knockdown as compared to siControl while *Hal*, *Efcab5* and *H2afy2* silencing did not alter efferocytosis (Figure 6A). We confirmed the engulfment of PKH67-labelled apoptotic Jurkat cells in si*Stard13* and si*Arsg*, but not si*Hal*, si*Efcab5* and si*H2afy*-treated BMDM by confocal microscopy (Figure 6B and Supplemental Figure 7B). We found significant decreases in *MerTk, Tyro3, Axl*, *Lrp1,* and *Rac1*, which are important genes in efferocytosis(57), and an increase in *RhoA*, which regulates actin dynamics(58), in BMDM after *Stard13* and *Arsg* downregulation (Figure 6C). In line with these data, we found a significant increase in the expression of *RhoA* and decreased *Rac* levels in *Rbpj*^-/-^ BMDM (Figure 6D). We also performed flow cytometry and immunofluorescence microscopy for MERTK, which is an efferocytosis receptor that recognizes and internalizes apoptotic cells by binding to phosphatidylserine on their surface (39). These experiments demonstrated that MerTK expression was significantly lower in BMDM after *Stard13* and *Arsg* silencing (Supplemental Figure 7C-D). Since ADAM17 cleaves MerTk(59, 60), we sought to examine if *Stard13* and *Arsg* controls this protease. We observed that *Adam17* expression was upregulated after *Stard13* or *Arsg* silencing, suggesting that reduced MerTK expression after *Stard13* and *Arsg* silencing is mediated by Adam17 (Figure 6E). Additionally, we observed decreased MerTK expression and elevated levels of ADAM17 in plaque macrophages of *LysM*^Cre/+^ *Rbpj*^fl/fl^ mice compared to *LysM*^+/+^ *Rbpj*^fl/fl^ mice, which is in line with the heightened necrotic core areas in atherosclerotic plaques of *LysM*^Cre/+^ *Rbpj*^fl/fl^ mice (Figure 6F). Consistently, BMDMs lacking *Rbpj* had reduced *Adam17* expression (Supplemental Figure 7E). We assessed pro- and anti-inflammatory genes after silencing *Stard13* and *Arsg* in BMDM (Supplemental Figure 7F). We found a significant upregulation of *Il1b* after *Stard13* silencing in BMDM whereas *Arsg* silencing mediated a non-significant increase in *Il1b* in these macrophages. In contrast, the anti-inflammatory genes *Il10* and *Tgfb* were downregulated in both groups. We next silenced *Stard13* and *Arsg in vivo* in peritoneal macrophages before injecting apoptotic Jurkat cells intraperitoneally to assess the impact of these genes in resident macrophage-mediated efferocytosis (Figure 6G and Supplemental Figure 7G). This resulted in significant reduction of apoptotic cell uptake by peritoneal macrophages (Figures 6H and 6I). Congruently, there was a decrease in the expression of the efferocytotic signaling genes and *Rac1* (Figure 6J). Conversely, *RhoA* expression heightened upon *in vivo* silencing of *Stard13* and *Ars*g. In aggregate, these findings demonstrate that *Stard13* and *Arsg* are crucial to macrophage-mediated efferocytosis.

Given our findings in mouse macrophages, we next sought to determine whether RBPJ similarly regulates STARD13 and ARSG to control efferocytosis in human macrophages. At first, we used Broad Institute Single Cell Portal to evaluate the expression and correlation of RBPJ and its downstream and upstream genes in publicly available single nuclei RNA sequencing data set of human carotid tissue(61). In the dataset we found increased expression of *NOTCH2*, *RBPJ*, *MERTK*, *STARD13*, and *ARSG* in plaque macrophages (Supplemental Figure 7H). We also found that the efferocytotic gene *MERTK* expression is correlated with *NOTCH2, RBPJ*, and *STARD13* (Supplemental Figure 7I). To validate whether RBPJ regulates efferocytosis in human macrophages, we used human monocyte-derived macrophages. *RBPJ* silencing in human monocyte-derived macrophages reduced efferocytosis of apoptotic Jurkat cells (Figure 7A-C). To test if RBPJ mediates efferocytosis through *STARD13* and *ARSG*, we overexpressed *STARD13 and ARSG,* in THIP-1 cells after *RBPJ* silencing with shRNA using a lentiviral vector and differentiated them into macrophages (Figure 7D-G). Overexpression of these genes abrogated suppressed efferocytosis in the macrophages lacking *RBPJ*, indicating that RBPJ mediates efferocytosis via *STARD13* and *ARSG*.

**Figure 7:**
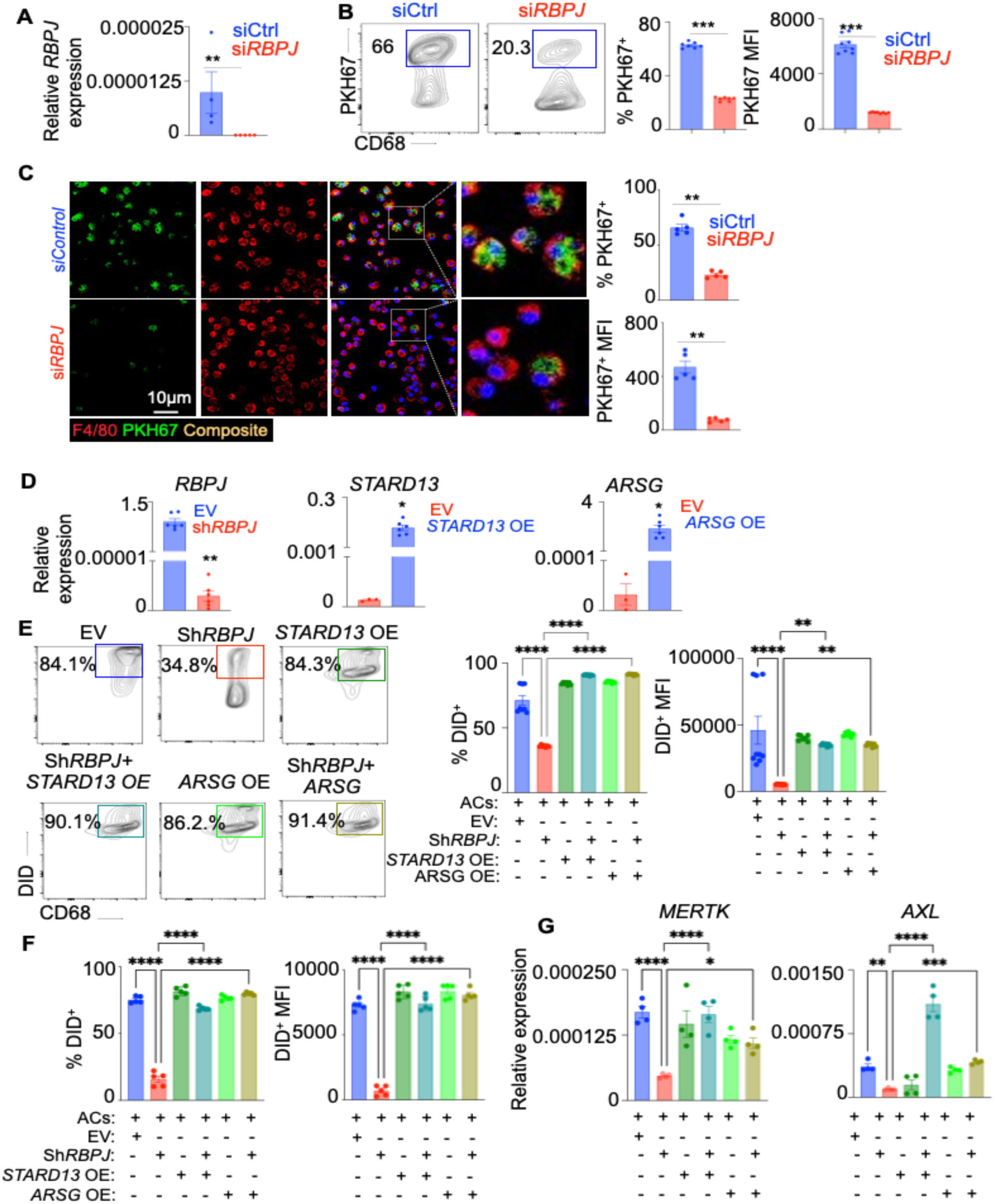
STARD13 and ARSG overexpression in human macrophages reverses impaired efferocytosis in the absence of *RBPJ*. **A-C.** Human monocyte-derived macrophages (hMDM) were treated with either si*Control* or si*RBPJ* and were co-cultured with PKH67^+^ ACs. **(A)** qPCR quantification of RBPJ expression (n=4–5/group). **B-C.** Efferocytosis was quantified by flow cytometry (n=6–7/group) (**B**) and confocal microscopy (n=5/group) (**C**). **D-G.** THP-1 cells were transduced with lentiviruses expressing empty vector (EV), sh*RBPJ*, *STARD13* (*STARD13* OE), or *ARSG* (*ARSG* OE) and differentiated into macrophages. These macrophages were co-cultured with DiD-labeled apoptotic Jurkat cells (ACs). **(D**) qPCR validation of RBPJ silencing and STARD13/ARSG overexpression (n=5-6/group). (**E**) Representative flow cytometry plot and quantification of efferocytosis (n=5/group). (**F**) Confocal microscopy-based assessment of efferocytosis (n=5–7/group). (**G**) qPCR quantification of *MERTK* and *AXL* expression in the THP-1 cells (n=4/group). The data are expressed as mean ± SEM and obtained from 3–5 independent experiments. \**p* < 0.05, \*\**p* < 0.01, \*\*\**p* < 0.001, \*\*\*\**p* < 0.0001.

### Stard13 sustains efferocytosis by controlling Rho and RAC GTPases balance and encouraging actin polymerization

During phagocytic clearance, macrophages internalize apoptotic cells by forming filopodia in an actin-dependent manner. The macrophage filopodia or phagocytic cup are rich in filamentous F-actin(62). Actin rearrangement during efferocytosis is regulated by the Rho family of small guanosine triphosphatase (GTPase)(63). To discern the mechanisms of Stard13/Arsg-facilitated efferocytosis, we used Rhosin, an inhibitor of Rho, which is known to mediate actin depolymerization. As expected, siRNA against *Stard13* and *Arsg* reduced apoptotic cell engulfment by macrophages (Figures 8A and 8B). Interestingly, Rhosin treatment restored efferocytotic ability of BMDM after *Stard13* silencing (Figure 8A). Surprisingly, we did not see similar effect in BMDM treated with Rhosin after *Arsg* downregulation (Figure 8B). In line with this finding, we observed an increase in RhoA and Cdc42 activation, and Rac1 inactivation in absence of *Stard13* (Figure 8C) and *Rbpj* (Supplemental Figure 8A) while we did not see any change in these GTPases in absence of *Arsg* (Figure 8C), suggesting that RBPJ-Stard13-mediated efferocytosis is dependent on Rho. To validate that Rho negatively regulates efferocytosis in absence of RBPJ, we treated *Rbpj*^+/+^ and *Rbpj*^-/-^ BMDM (Figure 8D) and si*Contro*l and si*Rbpj*-transfected BMDM (Supplemental Figure 8B) with Rhosin. Rho inhibition reversed suppressed efferocytosis in absence of RBPJ. We evaluated pro- and anti-inflammatory genes in macrophages after Rhosin treatment. We found that the pro-inflammatory gene *Il1b* was downregulated and the anti-inflammatory gene *Tgfb* was upregulated in si*Stard13* BMDM treated with Rhosin compared to si*Strard13*-treated BMDM without Rhosin treatment (Supplemental Figure 8C). Since Rho regulates actin polymerization, we quantified F/G-actin ratio in the absence of *Stard13* and *Arsg* in macrophages undergoing efferocytosis and found F/G ratio to be significantly decreased after si*Stard13* treatment and in *Rbpj* KO but not after si*Arsg* treatment (Figure 8E and Supplemental Figure 8D-E) suggesting that Stard13-mediated efferocytosis depends on F-actin polymerization. ARSG is a lysosomal resident protein that removes 3-O-sulfate from glucosamine and aids in lysosomal degradation(64). As such, we investigated ARSG expression in the lysosomal compartment during efferocytosis. Presence of apoptotic cells heightened ARSG signal in BMDM lysosomes, which was abrogated when *Rbpj* was absent (Figure 8F). Subsequently, we quantified the expression of the genes involved in lysosomal function and biogenesis(65) during efferocytosis in the presence or absence of RBPJ. We did not find a change in the expression of any of these genes (Supplemental Figure 8F).

**Figure 8:**
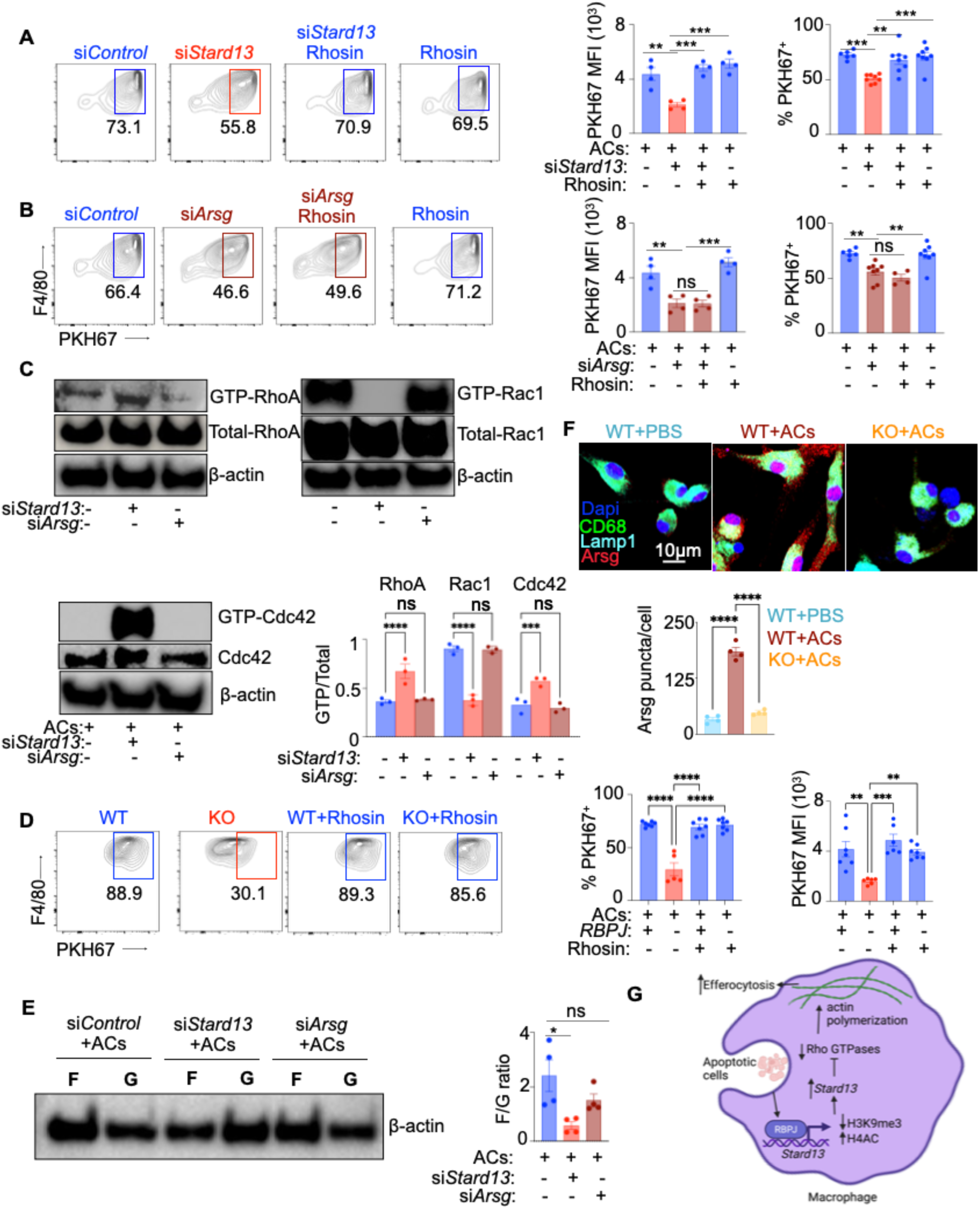
Stard13 mediates efferocytosis via Rho activation and actin polymerization. **A.** Representative flow cytometry plots and bar graphs showing PKH67^+^ apoptotic cell uptake by BMDM in presence or absence of Rhosin after si*Control* or si*Stard13* treatment. (n=5-7/group). **B.** Representative flow cytometry plots and bar graphs showing PKH67^+^ apoptotic cell uptake by BMDM in presence or absence of Rhosin after si*Control* or si*Arsg* treatment. (n=5-7/group). **C.** Immunoblot images and quantification of GTP or Total-RhoA, GTP or Total-Rac1 and GTP or Total-Cdc42 in BMDM cultured with apoptotic cells after silencing *Stard13* or *Arsg*. (n=5-7/group). **D.** Representative flow cytometry plots and bar graphs showing PKH67^+^ apoptotic cell uptake by *Rbpj*^+/+^ and *Rbpj*^-/-^ BMDM in presence or absence of Rhosin (n=5-7/group). **E.** Immunoblot images and quantification of F-actin and G-actin in BMDM cultured with apoptotic cells after silencing *Stard13* or *Arsg*. (n=5-7/group). **F.** Representative immunostaining images and quantification of Arsg in lysosomes of BMDM obtained from RBPJ^+/+^ (WT) or RBPJ^-/-^ (KO) mice. (n=5/group). **G.** Schematic diagram showing the summary of the study. The data are expressed as mean ± SEM and obtained from 3–4 independent experiments. \**p* < 0.05, \*\**p* < 0.01, \*\*\**p* < 0.001, \*\*\*\**p* < 0.0001.

## Discussion

We conclude that RBPJ facilitates phagocytotic clearance of apoptotic cells by activating *Stard13* by suppressing H3K9me3 mark on the *Stard13* promoter. Although endothelial RBPJ signaling is able to promote inflammatory leucocyte recruitment in atherosclerosis(30), we did not observe a role of myeloid RBPJ in leukocyte influx. Stard13 inactivates RhoA, promoting F-actin polymerization and apoptotic cell engulfment by macrophages (Figure 8G). Defective efferocytosis results in many diseases, including atherosclerosis, diabetes, and myocardial infarction(2, 39, 66). Thus, promoting the clearance of apoptotic cells will likely resolve inflammation and thwart the progression of chronic diseases. Our study suggests that this can be done in several ways: a) stimulating the Notch signaling pathway, b) increasing the expression of Stard13 and Arsg, c) epigenetic modifications of the promoters of these genes, and d) tailoring Rho GTPases. Future studies are warranted to ascertain the therapeutic benefits of modulating these pathways to promote efferocytosis, dampen inflammation, and mitigate disease pathogenesis.

As reported, RBPJ is known to exert its effects via the Notch signaling pathway, which involves cleavage of the Notch receptor by extracellular ADAM metalloproteases and an intracellular γ-secretase upon receptor ligand interaction(41, 42). Moreover, it is reported that the Notch-RBPJ pathway is involved in fibrous cap formation in atherosclerosis(67). Consistent with these observations, we found a decrease in efferocytosis when the Notch signaling was inhibited in the absence of RBPJ or with an inhibitor of γ-secretase. If the other components of the Notch signaling, such as Mastermind-like 1, are also involved in apoptotic cell clearance needs to be evaluated.

Histone modifications are essential to cellular processes ranging from gene expression to immune response. Various epigenetic writers and readers, mediating different epigenetic marks, have been reported to govern immune cell functions(9, 13, 68, 69). RBPJ has been shown to regulate downstream genes via acting as a transcriptional activator or repressor(22). Through genome-wide ChIP sequencing, multiple genomic sites, including promoter sites and enhancer sites, have been identified to interact with RBPJ under different physiological and pathological conditions(19). As a corepressor complex, for example, RBPJ recruits histone demethylase like Kdm5a/Lid(70–72) and Kdm1a/Lsd1(70, 73–75). Kdm5a/Lid directly binds with RBPJ, demethylates H3K4me3, and downregulates the expression of Notch target genes(71), while Kdm1a/Lsd1 indirectly binds with RBPJ via L3mbtl3(74). The trimeric complex, formed by RBPJ, MAML1 and NICD, is able to recruit the histone acetyltransferase Kat3b/Ep300 to Notch target genes, and upregulate the expression of genes via histone acetylation(76–78). In our study, RBPJ appears as a key epigenetic mediator of macrophage-mediated efferocytosis by altering H3K9me3, which is a repressive epigenetic mark.

Based on the RNA sequencing data, we have identified the RBPJ-dependent genes, especially *Stard13* and *Arsg*, as the effectors of RBPJ-controlled efferocytosis. *Stard13* is found to interact with Rho GTPases and is involved in the cell cytoskeleton rearrangement(53). Arsg closely associates with organelle membranes, especially the lysosomes, and the deficiency of Arsg could lead to mucopolysaccharidoses(64, 79). We observed that lysosomal expression of *Arsg,* an arylsufatase (80), elevated in efferocytosis. It remains to be deciphered how lysosomal heparan sulfate degradation affects efferocytosis. Additionally, the Cluster 4 genes seem to regulate efferocytosis negatively as they are shown to be downregulated in macrophages in presence of apoptotic cells. However, it is beyond the scope of this manuscript to specifically test the contribution of each of the Cluster 4 genes in RBPJ-dependent efferocytosis.

The Rho family of GTPase activation induces cytoskeletal rearrangement(81). RhoA, Rac1, and Cdc42 are the three most well-known members of the Ras superfamily of Rho GTPases(82, 83). Guanine nucleotide exchange factors (GEFs), GTPase-activating proteins (GAPs), and guanine nucleotide dissociation inhibitors (GDIs) are three groups of proteins that control their activity. By encouraging Rho GTPase’s intrinsic GTPase activity and assisting in the production of the inactive form, GAPs negatively regulate Rho GTPases(84). Recently, Stard13 has been shown to be a GAP for RhoA and Cdc42(53). Studies have confirmed that through its varied regulation of Rho GTPases, Stard13 performs unique roles in the migration and invasion of cancer cells(85–87). However, whether Stard13/Rho GTPases signaling is also involved in the process of efferocytosis is unknown. RhoA regulates the development of stress fibers and focal adhesions, both of which have an impact on cell contractility(88). Rac1 controls the development of lamellipodia, which is characterized by substantial actin polymerization at the cell’s moving edge driving cell membrane protrusion(89). Consistent with these reports, we see an increase in RhoA and a decrease in Rac1 activation after the silencing of *Stard13*, which promotes efferocytosis. In contrast to the studies cited above and our results, an article reported that RhoA inhibition by Stard13 leads to inhibition of actin stress fiber assembly(87). In line with our findings, published reports suggest that apoptotic cell engulfment is mediated by Rac1 and its upstream activators, while RhoA and its downstream effector-Rho kinase (ROCK) have inhibitory effects(90, 91). In contrary, it has been observed that there is less Rac1 activity in the phagocytic membrane, concomitant with F-actin disintegration and phagocytic cup closure (92, 93). The possible explanations behind these contradictions could be that efferocytosis and phagocytosis in macrophages lead to discrete molecular signaling(1–3).

In summary, we characterized how an epigenetic modification can control macrophage-mediated efferocytosis and demonstrated a novel role of RBPJ in facilitating this process. Since defective clearance of apoptotic cells is a hallmark of different acute and chronic diseases, these findings may have implications for future macrophage-based immunotherapeutic strategies. Our study has identified different proteins, such as RBPJ, Stard13, and Rho GTPases, which can be targeted for this purpose.

## Methods

### Mice

Ten-twelve weeks old C57BL/6 (#000664), *LyzM-cre* (#004781)*, Ldlr^-/-^* (#002207) and *Apoe^-/-^* (#002052) male and female mice were procured from the Jackson Laboratory. *Rbpj^fl/fl^*mice were obtained from RIKEN, Japan (#RBRC01071). We fed *Ldlr^-/-^* and *Apoe^-/-^* mice with a western diet (Envigo RMC Inc, #TD88137) as atherosclerotic mouse strains, to induce atherosclerosis. With bone marrow from either *LyzM^+/+^ Rbpj^fl/fl^* or *LyzM^cre/+^ Rbpj^fl/fl^* mice, we reconstituted *Ldlr^-/-^* mice. During all the procedures, mice were anesthetized with isoflurane (2–3%). As advised by the American Veterinary Association, carbon dioxide was used to euthanize the mice.

### Study approval

The Institutional Animal Care and Use Committee (IACUC) at the University of Pittsburgh gave its approval to conduct all the mouse experiments related to this report. Patients undergoing carotid endarterectomy were consented, and excised carotid artery plaques were collected. The University of Pittsburgh’s Institutional Review Board approved the human studies performed in this project (PRO18060512 and STUDY19020234).

### DOTAP mediated gene silencing in mice

For each mouse weighing around 22-25g, 33 µg of DOTAP (Encapsula NanoSciences, GEN-700) was used to deliver siRNAs (si*Rbpj,* si*Stard13* and si*Arsg*) i.p. Briefly, siRNA and DOTAP at a ratio of 1:7.5 were mixed, and the mixture volume was adjusted to 100 µl by adding RNAase and DNAase-free PBS. The mixture was incubated in ice for 15-20 mins and injected i.p in mice. Post 48 hours of siRNA delivery, the mice were euthanized and used for experiments(94).

### Gamma irradiation and bone marrow transplantation

Using a gamma irradiator, 10 Gy of radiation was applied to non-anesthetized *Ldlr*^-/-^ mice. The mice were then brought back to the animal facility and injected with one million bone marrow cells from either *LyzM^+/+^ Rbpj^fl/fl^* or *LyzM^cre/+^ Rbpj^fl/fl^* mice i.v. under anesthesia.

### Cell Culture

Human T lymphocytes cell line-Jurkat cells and mouse fibroblasts cell line-L929 were cultured in low glucose DMEM medium (Gibco, 11885-092) supplemented with 10% fetal bovine serum FB Essence (VWR, 10803-034), 100 μg/mL streptomycin, and 100 IU/ml penicillin (Gibco, 15140122). Cells were maintained in a humidified incubator with 5% CO2 and 37^0^C cell culture conditions.

### Isolation, culture and treatment of murine bone marrow-derived macrophage (BMDM)

After euthanizing mice with isoflurane, their leg bones including tibias, fibulas, and femurs were removed. DMEM supplemented with 4.5 g/L glucose, 20% L-929 conditioned media, 10% FBS, and 1% penicillin/streptomycin was used to flush the bones using a 26 gauge needle. Five 100 mm dishes were used to plate the cell suspensions after they had been passed through a 70 µm filter followed by centrifugation at 500 g. Cells were cultured for 4 days, at which point old media were removed, and the cells were washed with warm PBS to remove non-adherent cells and debris and replenished with fresh media. Cells were differentiated for 7–10 days with media replacement every 2–3 days(95). Cells were then collected for various experiments. Prior to treatments, cells were plated in 6-well or 12-well plates at a confluency of 1x 10^6^ cells/well or 0.5x 10^6^ cells/well respectively. These cells were then treated differently for various experiments with 1. 0.5 mg/mL oxLDL (ThermoFisher Scientific, L34357) or nLDL (MyBioSource, MBS143113) for 24 hours at 37°C. (96) 2. DAPT (Thermo Scientific, J65864.MA), a γ-secretase inhibitor, at a concentration of 10 µM for 24 hours under cell culture conditions(43) 3. N2-(4-isopropylphenyl)-5-(3-methoxyphenoxy)quinazoline-2,4-diamine (Yhhu-3792) (MCE, HY-120782), a Notch signaling activator, at a concentration of 2.5 µM for 24 hours under cell culture conditions of (44) 4. Rho inhibitor, Rhosin (MilliporeSigma, 555460) at a concentration of 30 µM for 24 hours in serum-free medium under cell culture conditions(97) and 5. Chaetocin, a lysine-specific histone methyltransferase inhibitor specific for SU(VAR)3-9, (MilliporeSigma, C9492) at a concentration of 0.5 µM for 72 hours under cell culture conditions(98)

### SiRNA transfection *in vitro*

Small interfering RNAs (siRNAs) targeting *Stard13*, *Arsg*, *Hal*, *Efcab5*, *H2afy2*, *Rbpj*, *Suv39h1,* and *Suv39h2* were transfected into BMDMs at a concentration of 4 nM using Lipofectamine 3000 Transfection Reagent (Invitrogen, L3000008) in accordance with the manufacturer’s instructions for gene-silencing studies. The cells were incubated with siRNA for 48 hours. With RT-qPCR, the knockdown efficiency was assessed.

### Quantitative polymerase chain reaction

Utilizing the rNeasy RNA isolation kit (Qiagen, 74104), total mRNA was extracted. The high-capacity RNA to cDNA synthesis kit (Applied Biosystems, 4387406) was used to generate cDNA from 100 ng of mRNA. SYBR green (Applied Biosystems, A25742) were used for quantitative RT-PCR, and the results were represented as Ct values normalized to the housekeeping gene-β actin. The sequences of the primers are provided in Supplemental Table 1.

### Organ harvesting and flow cytometry

Using cardiac puncture or retroorbital bleeding procedure, peripheral blood was drawn and collected in a tube containing a solution of 50 mM EDTA (Invitrogen, AM9260G). RBC in blood was lysed using a commercially available RBC lysis buffer (BioLegend, 420301). The mice were then perfused with PBS injection through the left ventricle. The aorta in full length was extracted, minced, and processed into single-cell suspension in an enzymatic combination of 450 U/mL collagenase I (Gibco, 17100017), 125 U/mL collagenase XI (Sigma-Aldrich, C7657), 60 U/mL dNase I (Qiagen, 79254), and 60 U/mL hyaluronidase (Sigma-Aldrich, H35006) at 37°C for 1 hour. The digested aortic tissue was then passed through 40 μm cell strainers. The cells were washed and resuspended in FACS buffer (0.5% BSA in PBS). Cells were then stained with fluorescently labeled antibodies for flow cytometry analysis as reported previously(99, 100). Specifically, the leukocytes suspensions obtained from aorta digestion were stained using the following antibodies: Ly-6C (Clone AL-21) (BD Biosciences, 563011), F4/80 (Clone BM8) (Biolegend, 123114), CD115 (Clone AFS98) (Invitrogen, 46-1152-82), Ly-6G (Clone 1A8) (Biolegend, 127614), and CD11b (Clone M1/70) (BD Biosciences, 557657). A BD LSR Fortessa flow cytometer was used to acquire samples and record data. Data were analyzed using FlowJo X 10.4.1 (Tree Star Inc.). Cell enumeration was performed using a hemocytometer prior to acquisition in the flow cytometer. Briefly, the counting chamber was loaded with a sample volume of 10 μL. For each sample, average leukocyte count was calculated using a minimum two mm^2^ area. Leukocyte sub-populations were calculated by multiplying the frequency of each population obtained by flow cytometry and the total number of leukocytes obtained by hemocytometer-assisted counts. Myeloid cells (CD19^low^ CD11b^high^), monocytes (CD19^low^ CD11b^high^ CD115^high^Ly-6G^low^), Ly-6C^high^ monocytes (CD19^low^ CD11b^high^ CD115^high^Ly-6G^low^ Ly-6C^high^), and neutrophils (CD19^low^ CD11b^high^ CD115^low^Ly-6G^high^) were identified as we previously described(99, 100). Live and dead cells were distinguished using viability dyes. Single cells were identified using forward- and side-scatter (FSC-A vs. FSC-W, SSC-A vs. SSC-W).

### Cell apoptosis analysis

*Rbpj*^+/+^ and *Rbpj*^-/-^ BMDM and apoptotic Jurkat cells were stained with propidium iodide (PI) and Annexin V using an apoptosis detection kit (ThermoFisher, V13245), according to the manufacturer’s instructions. Cells were acquired using a A BD LSR Fortessa flow cytometer and data were analyzed using FlowJo X 10.4.1 (Tree Star Inc.)

### Mouse atherosclerotic lesion analysis

Aortic roots were harvested, submerged in cryomold containing OCT (Fisher Scientific, 23730571), and immediately frozen on dry ice in 2-methylbutane bath. To measure the size of the plaque, the necrotic core area, and the thickness of the fibrous cap, 5 µm thick sections of aortic roots were sliced using a cryostat and stained with Masson’s trichrome (Polysciences, 25088-1). A Nikon 90-I microscope was used to acquire images. For each plaque, the average of five measures of the thinnest fibrous caps was calculated to determine the fibrous cap thickness. By quantifying the total acellular regions within each plaque, necrotic cores were examined.

### Immunofluorescence staining

For immunofluorescence staining, 0.1% triton X-100 was used to permeabilize tissue sections for an hour. After blocking, mouse tissue sections were stained with anti-F4/80 (clone A3-1, Invitrogen, MAI-91124), cleaved caspase-3 (Abcam, ab13847), RBPJ (Abcam, ab240227), CD68, (eBioscience, 14-0681-82), H3K9me3 (Active motif, 39161), Arsg, (Bioorbyt, orb318995), and Lamp1 (Thermofisher, 14-1071-82) followed by staining with fluorochrome-conjugated secondary antibodies. The sections were stained and fixed with Vectashield mounting medium with DAPI (Vector Laboratories, H-1200). Confocal microscopy (Nikon A1 Spectral Confocal) was employed to capture images. ImageJ software was used for image analysis.

### Human plaque histology

Patients undergoing carotid endarterectomy were consented, and excised carotid artery plaques were collected. The plaques were separated from the healthy part of the artery, fixed, frozen in OCT compound on dry ice. Using a cryostat, 10 µm sections were prepared from the tissue. The tissue sections were stained with antibodies against CD68 (Invitrogen, 14-0688-82) and RBPJ (Abcam, ab240227), followed by counterstaining with DAPI using Vectashield mounting medium (Vector Laboratories, H-1200). A Nikon A1 confocal microscope was used to capture the pictures, and Image J was used to quantify RBPJ expression in macrophages.

### Measurement of lipids

The Wako Cholesterol E kit (FUJIFILM Wako Chemicals USA Corporation, 99302501) was used to measure plasma cholesterol as per the manufacturer’s instructions.

### *In situ* efferocytosis

As previously stated(39), an in situ efferocytosis experiment was conducted. Briefly, to label lesional macrophages, sections of aortic root were stained with caspase-3 (Abcam, ab13847) followed by F4/80 (clone A3-1, Invitrogen, MAI-91124). After that, apoptotic cells were classified as either free (not associated with macrophages) or macrophage-related (colocalized or juxtaposed with macrophages). To illustrate the efficiency of efferocytosis, the results were presented as ratios of associated to free cells. A Nikon 90-I microscope was used to acquire images, and an observer who was blind to the samples’ group assignments used ImageJ to analyze the results.

### *In vitro* efferocytosis

BMDM (1×10^6^/well) were seeded in a 12-well plate and left overnight to adhere in cell culture conditions. Jurkat cells were tagged with PKH67 (Sigma-Aldrich, MIDI67-1KT) in accordance with the manufacturer’s instructions. Jurkat cells (3×10^6^) were then resuspended per ml of complete media and subjected to UV light exposure (254 nm, UVP) for 5 minutes. Following UV irradiation, cells were kept at 37°C and 5% CO_2_ for an hour. Using this method, we collected 80% of early apoptotic cells (Annexin V^+^ PI^−^). The Jurkat cells were then incubated with BMDM in a ratio of 5:1. After 45 minutes of incubation at 37°C, the medium was removed, and the macrophages were subjected to two cold PBS washes. To calculate the proportion of macrophages containing PKH67-tagged apoptotic cells, images were captured using a Nikon fluorescence microscope and analyzed using ImageJ.

### Ex vivo experiments for efferocytosis and apoptotic cell binding

Cells from the mouse peritoneal lavage were extracted and plated on 24-well plates. After two hours, the cells adhered to the plate. At this time, the medium was aspirated, and the cells were rinsed. F4/80^+^ macrophages made up more than 90% of the remaining adhering cells. Efferocytosis was performed as previously mentioned. After adding fluorescently tagged apoptotic Jurkat cells, plates were incubated at 4°C for an hour to assess the binding capability of macrophages. Following a cold PBS wash, plates were imaged as previously mentioned.

### Zymosan-induced peritonitis

One mg of zymosan A (Sigma-Aldrich, Z4250) resuspended in 500 μL sterile PBS (per mouse) was injected i.p. to induce peritonitis in mice. Mice were sacrificed after 6 hours, and exudates from the peritoneum were collected with 5 mL of sterile PBS.

### Peritoneal uptake of labeled aCs

The neutrophils from donor mice were removed by peritoneal lavage six hours after 1 mg zymosan A injection. The EasySep Mouse Neutrophil Enrichment Kit (Stemcell-Technologies, 19762A) was used to isolate and enrich the neutrophils from the collected lavage fluid as per manufacturer’s protocol. The extracted neutrophils were subsequently cultured for an overnight period in DMEM medium (Gibco, 11885092) containing 1% FB Essence (VWR, 10803-034) and 1% penicillin/streptomycin (Gibco, 15140122). The apoptotic neutrophils were tagged with PKH67 green the following day in accordance with the manufacturer’s instructions. The recipient mice were injected with 4×10^6^ of these tagged apoptotic neutrophils i.p. The mice were euthanized 45 minutes after the injection, and the peritoneum exudate was collected. The percentage of F4/80^+^ macrophages that had taken up one or more PKH67-labeled apoptotic cells was quantified to measure the extent of Efferocytosis (59)

### Acute lung injury

The mice were injected intratracheally with LPS (Sigma-Aldrich, L65291MG) at a dose of 3.75 μg/g body weight to induce acute lung injury. Mice were euthanized four days after LPS intratracheal injection. The blood was removed by cardiac puncture, and then the vasculature was perfused with cold PBS. The bronchoalveolar lavage (BAL) was collected after PBS injection into the lungs through the trachea. After this, the lungs were fixed by gravity perfusion of formalin.

### Lung immunofluorescence staining

Lung tissues that had been formalin-fixed were OCT-embedded and then sectioned. The lung tissue sections were stained with cleaved caspase-3 (Abcam, ab13847) followed by staining with a fluorochrome conjugated secondary antibody. The stained tissue sections were examined and quantitatively analyzed as described above.

### Cell sorting and RNA-seq analysis

The single cell suspensions of peritoneal macrophages were stained as described above, and the macrophages were sorted directly into RNA extraction buffer (PicoPure RNA Isolation Kit, Applied Biosystems, KIT0204) using a FACSAria III (BD Biosciences). The mRNA was extracted according to the manufacturer’s protocol. The mRNA sequencing was performed at Health Science Sequencing Core facility of UPMC Children’s Hospital of Pittsburgh. The acquired data were deposited to GEO (GSE207263). For performing downstream analysis, the FASTQ files were obtained and imported into CLC Genomics Workbench 21. A QC report was generated to check the quality of the data. The adaptor sequences (CTGTCTTATA) were trimmed from the 3’ end of the reads. To map the trimmed reads the *Mus musculus* CLC reference genome version 86 was used. The reads per kilobase of transcript per million mapped reads (RPKM) was used to calculate the gene expression. The differentially expressed genes (DEGs) among the groups were identified by keeping a cut off threshold of 2-fold change and FDR *P*< 0.05. The CLC Genomics Workbench 21 was used to represent data as PCA plots, volcano plots, and heatmaps.

### Ingenuity Pathway Analysis (IPA)

Pathway analysis of DEGs derived from bulk RNA sequences of tissue-resident (non-inflammatory) and monocyte-derived (inflammatory) macrophages was performed using the Qiagen IPA system 32718989 (101). The threshold for canonical pathway analysis was determined by using the −log (P-value) >2, where Z-score >2 indicated significant activation and Z-score <−2 indicated significant inhibition. Consistency scores for molecular networks and regulator effects were computed; a high consistency score denotes accurate regulatory effects analysis results (102).

### Chromatin Immunoprecipitation (ChIP)

ChIP was performed as previously described(103, 104). Briefly, passage-matched control and *RBPJ* KO BMDMs were fixed with 1% PFA (ThermoFisher Scientific, 28908) for 10 min at room temperature. The chromatin fragments of 200-500 base pairs were obtained by sonicating cells with a Bioruptor Pico (Diagenode, B01060010). The fragmented chromatin was incubated with Protein G Dynabeads (Invitrogen, 10004D) and one of the following antibodies: H3K9me3 (2 mg; Millipore, 05-1242), H4Ac (2 mg; Millipore, 05-1355), rabbit IgG (Abcam, ab171870), or mouse IgG (Abcam, ab37355). Phenol-chloroform was used to extract genomic DNA from immunoprecipitated (IP) and non-immunoprecipitated (INPUT) samples. TRANSFAC (geneXplain) and UCSC Genome Browser were used to identify binding motifs on gene promoters for RBPJ, H3K9me3, and H4AC. NCBI Primer-BLAST was used to design the sequences of the primers flanking the binding motifs. qPCR was performed to quantify precipitated binding motifs. The results were expressed as IP/INPUT.

### CUT&RUN Assay

The chromatin profiling was performed using the Cut and Run assay kit (Cell signaling technology, 86652) as per the manufacturer’s protocol. Briefly, 100,000 BMDM plated in 12-well plates, were collected, in Eppendorf tubes, washed, and incubated with activated Concanavalin A beads for five minutes under rotating conditions. The supernatant was removed using a magnetic stand, and the cells were then resuspended in antibody binding buffer containing 1X Spermidine, 1X PIC, and 0.01% digitonin. To this 2:100 µl of anti-H3K9me3 antibody (2mg; Millipore, 05-1242) was added and rotated at 4^0^ C overnight. Next day, the cells were collected and treated with 1.5 µl of pA/G-mNase enzyme for 1 hour under rotating conditions at 4^0^ C to facilitate enzyme binding. Next, the cells were treated with 3 µl of cold CaCl_2_ and incubated at 4^0^ C for an hour to initiate enzyme activation for digestion and release of targeted chromatin. The genomic DNA was purified using DNA Purification Buffers and Spin Columns (Cell signaling technology, 14209) as per the kit’s instruction. Sequencing libraries were prepared using CUT&RUN Library Prep Kit (EpiCypher, SKU: 14-1001). We performed the sequencing of extracted genomic DNA with the help of Health Science Sequencing Core located at UPMC Children’s Hospital of Pittsburgh.

### CUT&RUN data-processing and analysis

The analysis of CUT&RUN DNA sequencing was performed by using the Partek Flow (version 10.0.22.1204) software. In brief, pre-alignment QA/QC was first performed and then adaptors were trimmed before BWA alignment to the Mus musculus GRCm39 (mm39) genome. Aligned reads were then checked for post-alignment QA/QC and filtered. Identification of broad H3K9me3 peaks was carried out using the MACS2 peak function. The peaks were annotated using Ensembl transcript release 106. The regions were quantified using 80% minimum read overlap with feature. The region counts obtained were then normalized and annotated. Two Gene Specific Analysis (GSA) was performed 1. WT+ACs vs. WT+PBS and 2. RBPJ KO +ACs vs. WT+ACs using with FDR step up= 0.05 and Log Fold change +/-2. The list was also filtered for all upregulated and downregulated regions upon which gene set enrichment (GSEA) analysis was performed. The GSEA data of CUT&RUN was integrated with Dseq2 data of Bulk RNA seq to obtain the common genes. The acquired data were deposited to GEO (GSE253834). The annotated peaks are provided in Supplemental Table 2.

### RhoA/Rac1/Cdc42 Pull down assay

*Rbpj*^+/+^ and *Rbpj^-/-^* BMDM and BMDM treated with *Stard13* and *Arsg* siRNAs undergoing efferocytosis were lysed. GTP-RhoA, GTP-Rac, and GTP-Cdc42 from the cell lysates were pulled down using RhoA/Rac1/Cdc42 Activation Assay Combo Kit (Cell BioLabs, NC0373752) as per the manufacturer’s protocol. Briefly, the cell lysates were gently shaken with GST-CRIB or GST-RBD for an hour at 4 °C. The samples were then centrifuged, and the resulting pellet were washed for 2-3 times. The detection of GTP-RhoA, GTP-Rac1, and GTP-Cdc42 was performed by immunoblotting using respective antibodies supplied with the kit. To detect the presence of total amount of RhoA, Rac1, Cdc42, and GAPDH proteins, whole cell lysates were used.

### F/G actin estimation

*Rbpj*^+/+^ and *Rbpj*^-/-^ BMDM (2×10^6^) and BMDM treated with *Stard13* and *Arsg* siRNAs were subjected to efferocytosis by co-culturing them with apoptotic Jurkat cells in a 60 mm petri dishes. These cells were washed and lysed with ice-cold PBS and actin stabilization buffer (0.1 M PIPES, pH 6.9, 30% glycerol, 5% DMSO, 1 mM MgSO4, 1 mM EGTA, 1% TX-100, 1 mM ATP, and protease inhibitor), respectively, for 10-12 mins on ice. The cell lysates were collected and centrifuged at 16,000xg at 4°C for 75 minutes. The pellets containing F-actin were resuspended with actin depolymerization buffer (0.1 M PIPES, pH 6.9, 1 mM MgSO4, 10 mM CaCl2, and 5 μM cytochalasin D), and the supernatants containing G-actin were collected in a separate microcentrifuge tube. A 12% SDS-PAGE gel was used to resolve the G and F actin fractions, which was further immunoblotted with anti β-actin (Milipore, A3854) antibody. Densitometric analysis of the immunoblot was performed using NIH Image J software, and the data were represented as F/G ratio.

### Statistics

Data are represented as mean±SEM. Statistical significance between groups was assessed, and graphs were created using Prism. Non-parametric t test was used to compare differences between two group, and the one-way ANOVA was used for data sets containing more than two groups. Results were considered as statistically significant when *P*<0.05.

## Data availability

The article and its online supplementary files contain all of the information that the authors claim to have.

## Author contributions

SS and XZ were involved in designing and executing the experiments, data analysis and manuscript writing. LL was involved in performing and analyzing the ChIP-qPCR experiments. JS assisted in mice organ harvest and IJ and GM helped with the data analysis. PD was involved in designing experiments and writing the manuscript and provided the fund to complete the study.

## Acknowledgments

This research was funded by the National Institutes of Health (NIH) grants R00HL121076, R01HL142629, R01HL142629-04W1, R01HL143967, R01AG069399 and R01DK129339, the American Heart Association (AHA) Transformational Project Award (19TPA34910142), the AHA Innovative Project Award (19IPLOI34760566), the ALA Innovation Project Award (IA-629694), and the AHA Innovative Project Award (23IPA1053549) to P. Dutta. S. Sadaf received funds from the AHA Postdoctoral Fellowship Award (23POST1029135), and X. Zhang has been awarded the Young Scientists Fund of the National Natural Science Foundation of China (No. 82100346 and 82370441). The Center for Biologic Imaging (CBI), University of Pittsburgh, used NIH-supported grants to perform confocal and intravital microscopy. Specifically, the NIH grant 1S10OD019973-01 was used to fund the confocal microscope. The bulk RNA seq data analysis was carried out using CLC Genomics Workbench software, which is licensed through the Molecular Biology Information Service of the Health Sciences Library System (HSLS), University of Pittsburgh. Partek Flow software, version 10.0.22.1204, licensed by the HSLS University of Pittsburgh, was used to analyze CUT&RUN sequencing data. The University of Pittsburgh Center for Research Computing supplied the resources necessary to support this research, in part. The BioRender software was used to create the visual abstract.

---

**Supplemental Figure 1:**
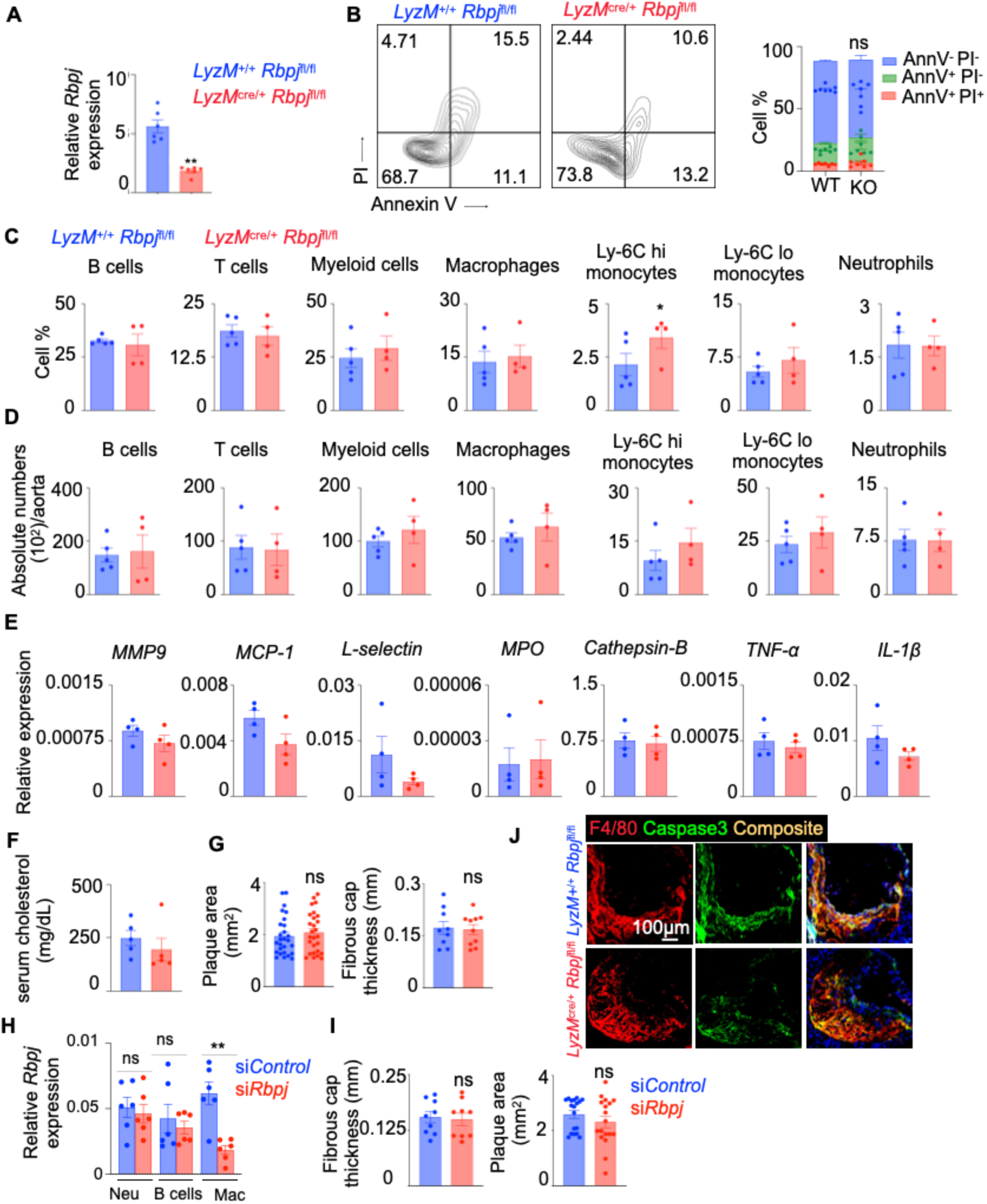
*RBPJ* deficiency does not alter inflammatory cell numbers, plaque areas, and fibrous cap thickness in mice with atherosclerosis. **A.** qPCR quantification of *Rbpj* in *LyzM^+/+^ Rbpj^fl/fl^* and *LyzM^cre/+^ Rbpj^fl/fl^* BMDM. (n=6/group). **B.** Representative flow plots showing Annexin V and PI staining in *LyzM^+/+^ Rbpj^fl/fl^* and *LyzM^cre/+^ Rbpj^fl/fl^* BMDM and the quantification of AnnV^+^ PI^+^ (dead), AnnV^+^ PI^-^ (apoptotic), and AnnV^-^ PI^-^ (live) cells. (n=7/group). **C-D.** Quantification of frequencies (C) and numbers (D) of different leukocyte subsets in the aortas of *Ldlr*^-/-^ *LyzM^+/+^ Rbpj^fl/fl^* and *Ldlr*^-/-^ *LyzM^cre/+^ Rbpj^fl/fl^* mice assessed by flow cytometry. (n=4-5/group). **E.** qPCR quantification of the inflammatory genes in the aortic arches of these mice. (n=4/group). **F.** Serum cholesterol levels of *Ldlr*^-/-^ *LyzM^+/+^ Rbpj^fl/fl^*and *Ldlr*^-/-^ *LyzM^cre/+^ Rbpj^fl/fl^* mice assessed by ELISA. (n=4-5/group). **G.** Quantification of plaque area and fibrous cap thickness using Masson’s trichome staining of aortic root sections of *Ldlr*^-/-^ *LyzM^+/+^ Rbpj^fl/fl^* and *Ldlr*^-/-^ *LyzM^cre/+^ Rbpj^fl/fl^* mice. (n=4-5/group). **H.** qPCR analysis of *Rbpj* in aortic neutrophils, B cells, and macrophages from HFD-fed *ApoE⁻/⁻* mice treated with either siControl or si*Rbpj* (n=6/group). **I.** Quantification of plaque area and fibrous cap thickness using Masson’s trichome staining of aortic root sections of HFD-fed *ApoE⁻/⁻* mice treated with either siControl or si*Rbpj* (n=6/group). **J.** Representative confocal images of aortic root sections of *Ldlr*^-/-^ *LyzM^+/+^ Rbpj^fl/fl^* and *Ldlr*^-/-^ *LyzM^cre/+^ Rbpj^fl/fl^* to detect apoptotic cells. (n=4-5/group). The data are expressed as mean ± SEM and obtained from 3 independent experiments. \*\**p* < 0.01.

**Supplemental Figure 2:**
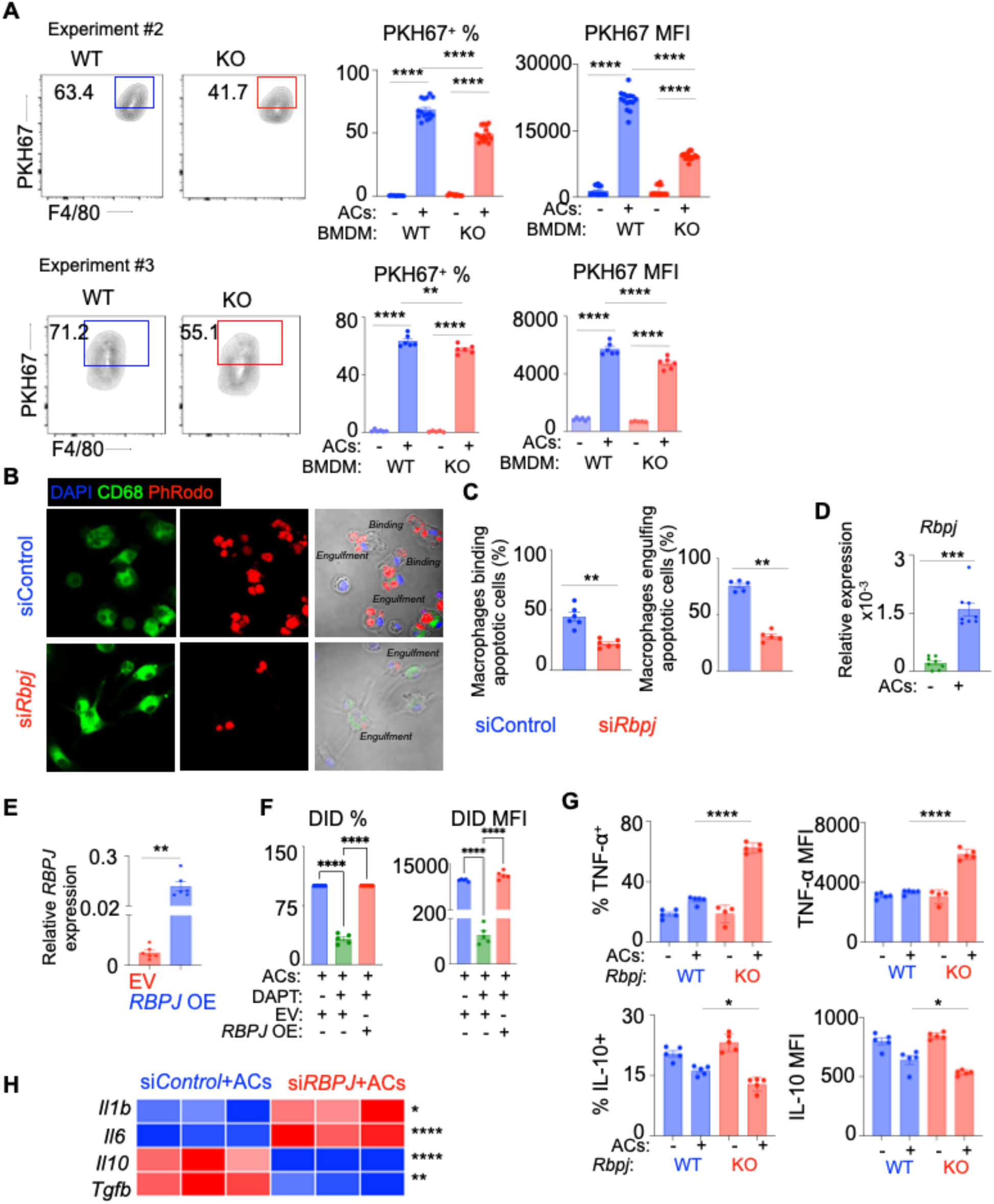
RBPJ deficiency impairs *in vitro* efferocytosis and augments inflammatory pathway in macrophages. **A**. Flow cytometry plots showing the uptake of pHrodo^+^ apoptotic Jurkat cells (ACs) by WT and *Rbpj* KO BMDM (n=6-8/group). **B-C**. Representative confocal images (**B**) and quantification (**C**) showing the frequencies of macrophages binding or engulfing ACs in si*Control* and si*RBPJ-*treated BMDM (n=5/group). **D.** qPCR data showing *Rbpj* expression in presence and absence of apoptotic Jurkat cells (n=8/group). **E.** qPCR analysis for *RBPJ* expression in empty vector control (EV) and *RBPJ*-overexpressing (*RBPJ* OE) THP-1 macrophages (n=6/group). **F.** Efferocytosis quantification of confocal images after DAPT treatment in THP-1 macrophages with or without *RBPJ* overexpression (n=5/group). **G.** The frequency of TNF-⍺^+^ and IL-10^+^ macrophages and the MFI of these cytokines were quantified by flow cytometry in WT and KO BMDM during efferocytosis (n=4-5/group). **H.** The heatmap showing qPCR quantification of *Il1b*, *Il6*, *Il10,* and *Tgfb* expression in si*Control* and si*RBPJ-*treated BMDM cultured with apoptotic cells. n=3-6/group. The data are expressed as mean ± SEM and obtained from 3 independent experiments. \**p* < 0.05, \*\**p* < 0.01, \*\**p* < 0.001, \*\*\*\**p* < 0.0001.

**Supplemental Figure 3:**
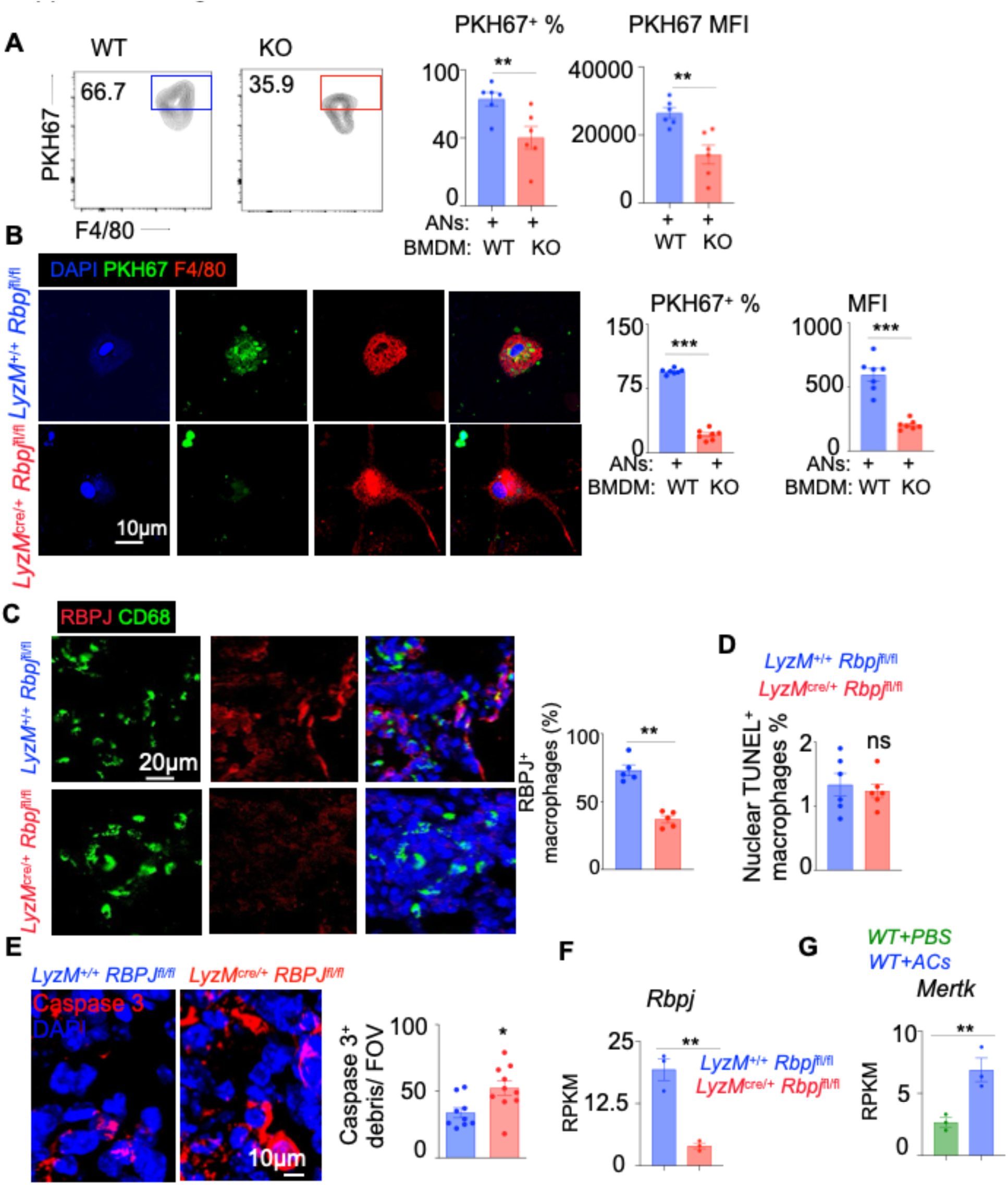
RBPJ deficiency impairs efferocytosis in mouse models of acute lung injury and peritonitis. **A-B.** Flow cytometry plots (**A**) and confocal images (**B**) showing the uptake of PKH67^+^ apoptotic neutrophils (ANs) by WT and *Rbpj* KO BMDM and the quantification of the frequency of PKH67^+^ BMDM and PKH67 MFI. (n=6-8/group). **C**. Lung sections of *LyzM*^+/+^ *Rbpj*^fl/fl^ (WT) and *LyzM^cre^*^/+^ *Rbpj*^fl/fl^ (KO) having acute lung injury (ALI) were stained for CD68 (green) and RBPJ (red). The frequencies of RBPJ^+^ macrophages were calculated (n=5/group). **D.** Nuclear TUNEL^+^ CD68^+^cells were quantified in lung sections of WT and KO mice with ALI (n=5/group). **E.** Representative confocal images and bar graphs showing caspase 3^+^ cells in the lungs. (9-10/group). **F-G.** Bulk RNA sequencing was performed in *Rbpj*^+/+^ (WT) and *Rbpj*^-/-^ (KO) peritoneal macrophages exposed or non-exposed to ACs (n=3/group). The RPKM values for *Rbpj* (**F**) in *Rbpj^+/+^*and *Rbpj^-/-^* peritoneal macrophages exposed to ACs and *Mertk* (**G**) in *Rbpj^+/+^* peritoneal macrophages cultured in the absence (WT+PBS) or presence (WT+ACs) of apoptotic Jurkat cells are shown. The data are expressed as mean ± SEM and obtained from 3 independent experiments \*\**p* < 0.01, \*\*\**p* < 0.001, \*\*\*\**p* < 0.0001.

**Supplemental Figure 4:**
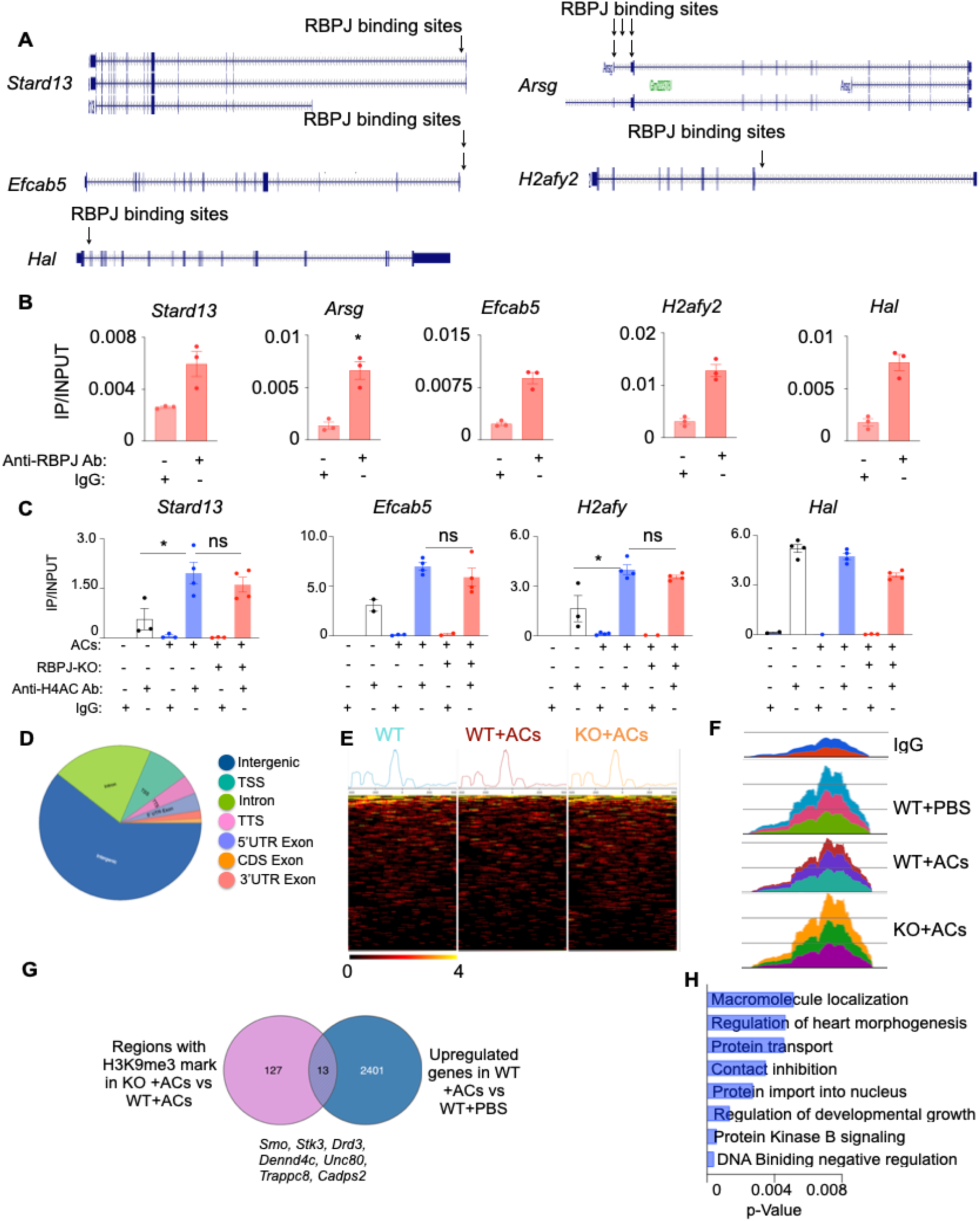
RBPJ binds to the promotor regions of the RBPJ-dependent genes, and its deficiency amplifies a genome wide H3K9me3 enrichment during efferocytosis. **A.** Binding sites of RBPJ in the *Stard13, Arsg, Efcab5, H2afy2,* and *Hal* promoter regions using the UCSC Genome Browser. **B.** ChIP-qPCR quantification of RBPJ occupancy on the *Stard13, Arsg, Efcab5, H2afy2,* and *Hal* promoters in BMDM in the presence of apoptotic Jurkat cells (n=3/group). The data from one of two independent experiments are shown. **C.** ChIP-qPCR quantification of H4AC mark in the promotor regions of *Stard13, Efcab5, H2afy2,* and *Hal* in the presence or absence of apoptotic Jurkat cells in BMDM of *LyzM^+/+^ Rbpj^fl/fl^* (WT) and *LyzM^cre/+^ Rbpj^fl/fl^* (KO) mice. (n=3-4/group). The data are obtained from 3 independent experiments. \**p* < 0.05. **D.** CUT&RUN analysis of genome-wide H3K9me3 distribution in BMDM obtained from *LyzM^+/+^ Rbpj^fl/fl^* (WT) and *LyzM^cre/+^ Rbpj^fl/fl^* (KO) in the presence (ACs) or absence (PBS) of apoptotic cells. (n=3/group). **E.** Heatmaps generated with the CUT&RUN data depicting H3K9me3 peaks enriched at regions spanning the transcription start sites (± 500 bp). (n=3/group). **F.** H3K9me3 distribution at random genomic regions for all the samples. n=3/group. **G.** Venn diagram indicating the 13 common genes in 1. H3K9me3 enriched regions of *RBPJ*^-/-^ vs. *RBPJ*^+/+^ BMDM cultured with apoptotic cells and 2. upregulated genes in response to apoptotic cells in *RBPJ*^+/+^ macrophages. **H.** Gene set enrichment analysis of the 13 common genes. The data are obtained from one independent experiment. The data are expressed as mean ± SEM. \**p* < 0.05.

**Supplemental Figure 5:**
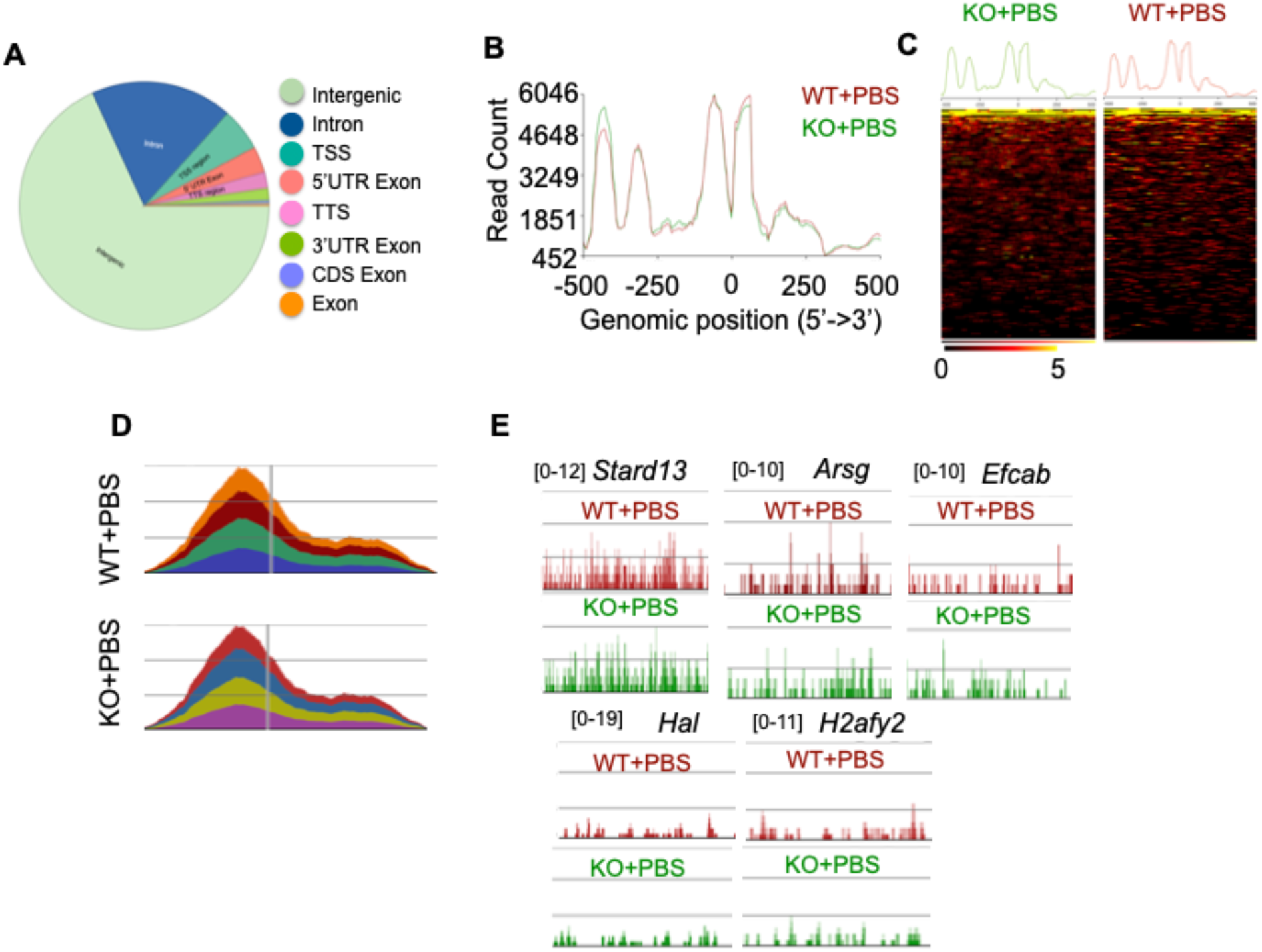
Myeloid *Rbpj* deficiency does not impact the H3K9me3 mark at baseline. CUT&RUN to detect H3K9me3 was performed in BMDM obtained from *LyzM*^+/+^ *Rbpj*^fl/fl^ (WT) and *LyzM*^cre/+^ *Rbpj*^fl/fl^ (KO) mice. The BMDM were cultured in the absence of apoptotic cells. **A.** A genome-wide H3K9me3 distribution is shown. **B.** The intensity plot of CUT&RUN analysis depicts H3K9me3 enrichment around ± 500 bp of the transcription start sites **C.** The heatmaps show the H3K9me3 peaks enriched at the regions spanning the transcription start sites (± 500 bp). **D.** H3K9me3 distribution at random genomic regions of WT and *Rbpj* KO BMDM. **E.** H3K9me3 peak distribution at the RBPJ-dependent genes *Stard13*, *Arsg*, *Efcab5*, *H2afy2*, and *Hal* is shown (n=4 per group).

**Supplemental Figure 6:**
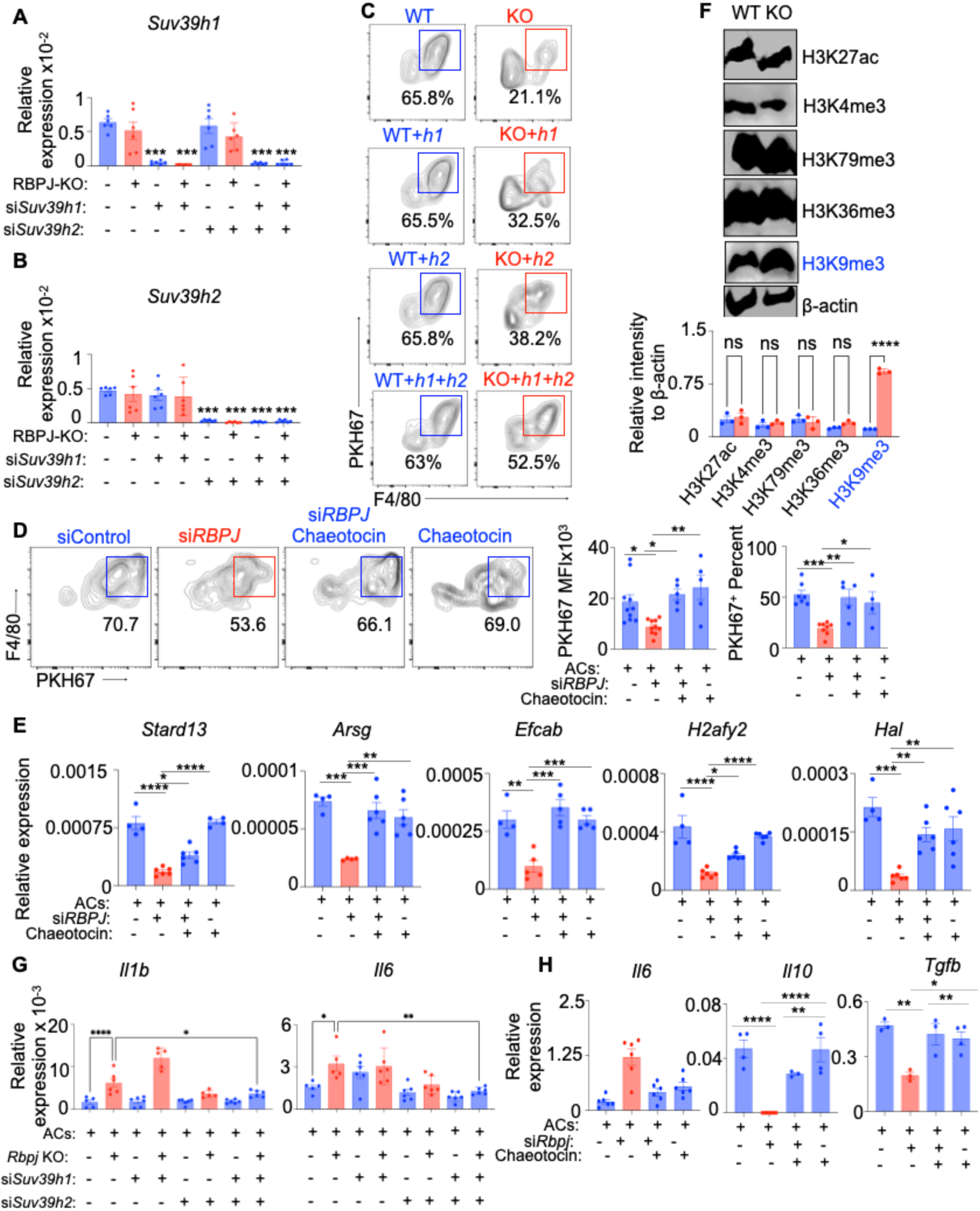
Decreased H3K9me3 expression drives RBPJ-mediated efferocytosis and augments the expression of the RBPJ-dependent genes. **A-B.** qPCR quantification of *Suv39h1* (**A**) and *Suv39h2* (**B**) in si*Control*, si*Suv39h1*, si*Suv39h2*, and si*Suv39h1/* si*Suv39h2-* transfected BMDM of *LyzM*^+/+^ *Rbpj*^fl/fl^ (WT) and *LyzM*^cre/+^ *Rbpj*^fl/fl^ (RBPJ KO) mice. (n=6/group). **C.** Representative flow cytometry plot showing PKH67^+^ apoptotic Jurkat cell uptake by the BMDM in A. (n=4-5/group). **D.** Representative flow cytometry plot and quantification of PKH67^+^ apoptotic Jurkat cell uptake by BMDM in presence or absence of si*Rbpj* and chaetocin. (n=5-7/group). **E.** qPCR quantification of the RBPJ-dependent genes: *Stard13, Arsg, Efcab5, H2afy2*, and *Hal* in BMDM cultured with apoptotic Jurkat cells in presence or absence of si*Rbpj* and chaetocin. (n=5-10 /group). **F.** Immunoblot images of H3K27Ac, H3K4me3, H3K79me3, H3K36me3, H3K9me3, and β-actin and their quantification relative to β-actin. (n=3/group) **G.** qPCR quantification of *Il1b* and *Il6* in the presence and absence of *Rbpj*, si*SUV39h1*, and si*SUV39h2* (n=5-10/group) **H**. qPCR quantification of *Il6, Il10, and Tgfb* in the presence or absence of si*Rbpj* and chaeotocin in BMDM cultured with apoptotic Jurkat cells (n=3-6/group). The data are expressed as mean ± SEM and obtained from 3 independent experiments. \**p* < 0.05, \*\**p* < 0.01, \*\*\**p* < 0.001, \*\*\*\**p* < 0.0001.

**Supplemental Figure 7:**
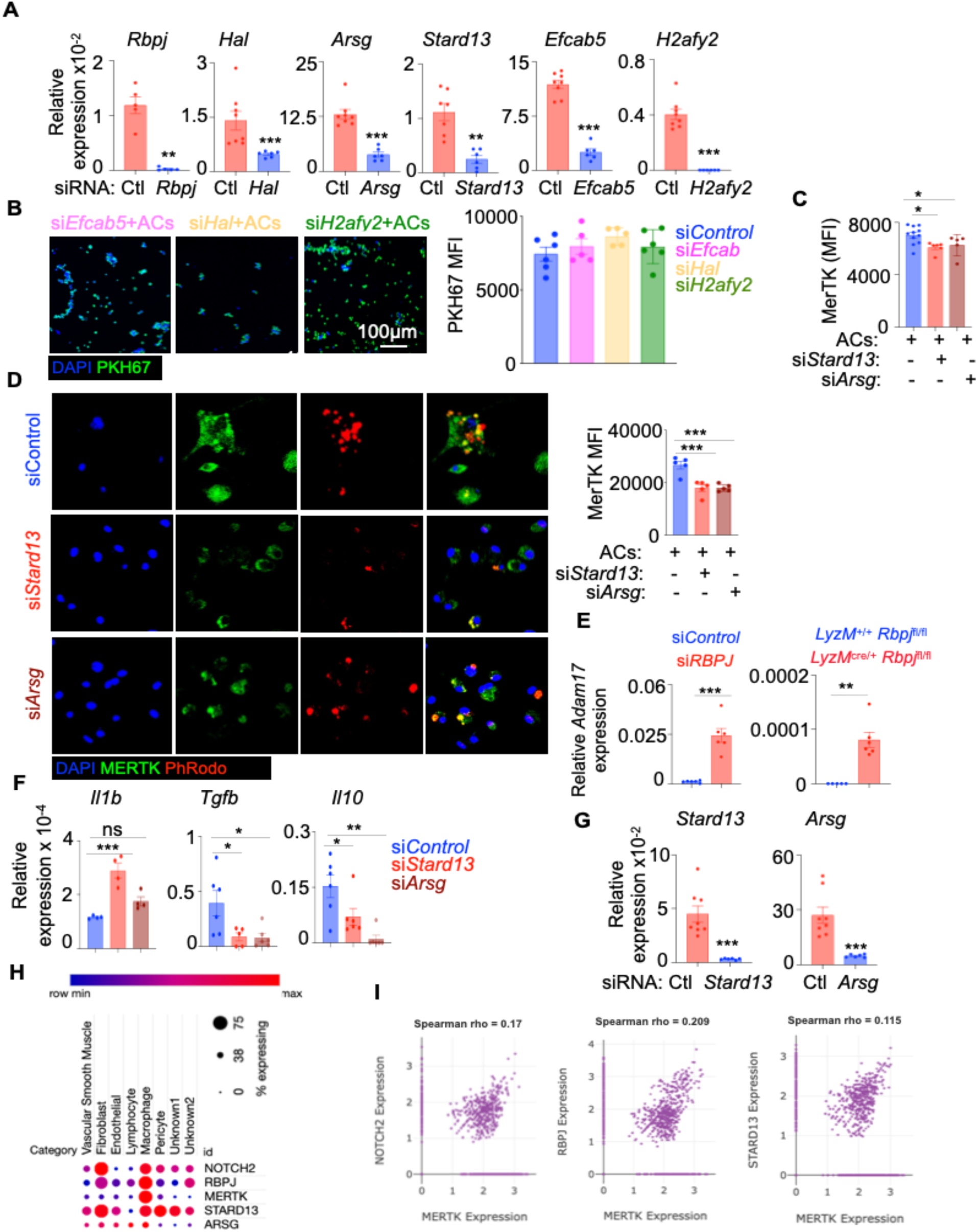
RBPJ-mediated efferocytosis is driven by Stard13 and Arsg. **A.** qPCR validation of *Rbpj*, *Hal*, *Arsg*, *Stard13*, *Efcab5,* and *H2afy2* silencing in BMDM. (n=5-8/group). **B.** Representative microscopy images and quantification of PKH67-labeled apoptotic Jurkat cell engulfment by BMDM after *Efcab5, Hal* or *H2afy2* silencing. (n=5/group). **C-D.** Flow cytometry quantification (**C**) (n=5-11/group) and confocal images and quantification (**D**) (n=5/group) showing MerTK expression in efferocytotic BMDM. **E** qPCR analysis of *Adam17* expression in BMDM transfected with either si*Control or siRBPJ,* or derived from WT and *Rbpj* KO mice (n=5-6/group). **F.** qPCR quantification of *Il1b, Tgfb,* and *Il10* in BMDM cultured with ACs in the presence or absence of si*Stard13* or si*Arsg* (n=5-6/group). **G.** qPCR validation of the silencing of *Stard13* and *Arsg* in peritoneal macrophages. (n=7-8/group). **H-I.** Single-nucleus RNA sequencing (snRNA-seq) data from human carotid atherosclerotic plaques were obtained from the Broad Institute Single Cell Portal. **H.** The bubble plot shows the expression of the genes (X-axis) in various cell populations (Y-axis). The dot size represents the percentage of cells expressing the genes, and the colors indicate the expression levels of the genes. **I.** The scatter plots display the correlations between *MERTK* and *NOTCH2*, *RBPJ*, and *STARD13* expression in plaque-resident macrophages. The data excluding the snRNA-seq data are expressed as mean ± SEM and obtained from 3 independent experiments. \*\**p* < 0.01, \*\*\**p* < 0.001, \*\*\*\**p* < 0.0001.

**Supplemental Figure 8:**
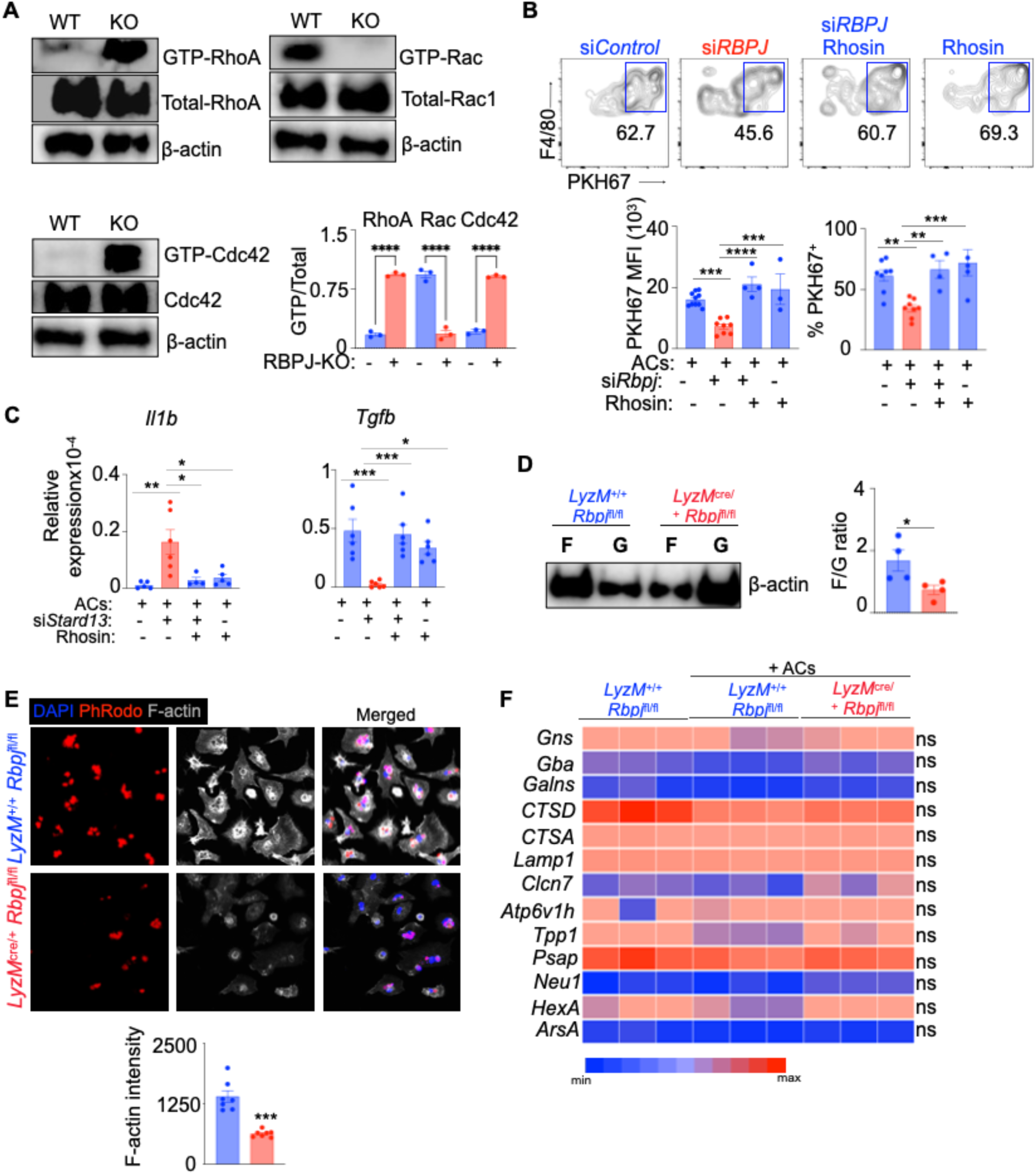
RBPJ-mediated efferocytosis is driven by Rho/Rac-driven actin polymerization guided by Stard13. **A.** Immunoblot images and quantification of GTP-RhoA, total-RhoA, GTP-Rac1, total-Rac1, GTP-Cdc42, and total-Cdc42 in *Rbpj*^+/+^ and *Rbpj*^-/-^ BMDM cultured with apoptotic cells. (n=5-7/group). **B.** Representative flow cytometry plots and bar graphs showing PKH67^+^ apoptotic cell uptake by BMDM in the presence or absence of Rhosin after si*Control* and si*Rbp*j transfection. (n=5-7/group). **C.** qPCR quantification of *Il1b* and *Tgfb* expression in BMDMs treated with si*Stard13* and Rhosin and co-cultured with apoptotic Jurkat cells. (n = 4–6/group).**D.** Immunoblot images and quantification of F-actin and G-actin in *Rbpj*^+/+^ and *Rbpj*^-/-^ BMDM cultured with apoptotic cells. (n=3/group). **E.** Representative confocal images and the bar graph depict F-actin intensity in *Rbpj*^+/+^ and *Rbpj*^-/-^ BMDM cultured in presence of PHrodo-labelled ACs (n=7/group). **F.** Heatmap of qPCR quantification of the genes crucial in the lysosomal degradation pathway in *LyzM^+/+^ Rbpj^fl/fl^* and *LyzM^cre/+^ Rbpj^fl/fl^* BMDM cultured in the presence or absence of apoptotic cells. (n=5/group). The data are expressed as mean ± SEM. \**p* < 0.05, \*\**p* < 0.01, \*\*\**p* < 0.001, \*\*\*\**p* < 0.0001.

**Supplemental Table 1:** This table lists the sequences of primers used for the qPCR and ChIP-qPCR experiments.

**Supplemental Table 2:** This table lists the annotated peaks of the CUT&RUN data.

**Supplemental Video 1:** BMDM derived from *LyzM^+/+^ Rbpj^fl/fl^* (WT) mice were co-cultured with apoptotic Jurkat cells (ACs) labelled with PHrodo. A timelapse imaging was performed to assess binding and engulfment of the Acs by the macrophages.

**Supplemental Video 2:** BMDM derived from *LyzM^cre/+^ Rbpj^fl/fl^* (KO) mice were co-cultured with apoptotic Jurkat cells (ACs) labelled with PHrodo. A timelapse imaging was performed to assess binding and engulfment of the Acs by the macrophages.

